# Phosphorylation of the α subunit inhibits proteasome assembly and regulates cell division in an archaeon

**DOI:** 10.1101/2025.05.09.653063

**Authors:** Ya Wu, Qi Gan, Kanghui Ning, Ran Zhang, Shikuan Liang, Yunfeng Yang, Pengju Wu, Xu Feng, Qunxin She, Jinfeng Ni, Yulong Shen, Qihong Huang

## Abstract

Archaea of the order Sulfolobales possess a eukaryotic-like cell division machinery and display a eukaryotic-like cell cycle, however, the cell division and cell cycle control mechanisms remain enigmatic. Here, we demonstrate that phosphorylation of the α subunit by a eukaryotic-like protein kinase ePK2 affects 26S proteasome assembly and controls cell division in *Saccharolobus islandicus*. ePK2 exhibits cell cycle-dependent expression at both transcriptional and translational levels. Deletion or overexpression of *ePK*2 results in impaired cytokinesis, with the deletion cells being unable to generate single chromosome cells after synchronization and the overexpression cells exhibiting growth retardation and cell enlargement. Interestingly, overexpression of ePK2 leads to a coherent reduction in cellular proteasome activity and degradation of cell division proteins. We identify S200 and T213 of the proteasome α subunit as specific target sites for ePK2 phosphorylation. Functional and structural analyses of site-directed mutants at S200 and T213 suggest that S200 phosphorylation disrupts the assembly of 20S into 26S proteasome whereas T213 phosphorylation interferes the *de novo* α ring assembly. Collectively, our study uncovers an ingenious and efficient mechanism of proteasome phosphorylation-mediated cell division regulation, a prototype of the eukaryotic cell cycle regulation system, in Sulfolobales archaea.

## Introduction

The eukaryotic cell cycle consists of four main phases, G1 (the first gap), S (DNA synthesis), G2 (the second gap), and M (mitosis) phases^1^. The cell cycle is strictly regulated by a combination of cyclins and cyclin-dependent kinases (CDKs)^1–3^. Accumulation of specific cyclins during different cell cycle stages is accomplished by cell cycle-dependent transcription and inhibition of proteasome-mediated degradation. In turn, this transcription is dependent on CDK-mediated phosphorylation and activation^4,5^. The proteasomes are regulated by phosphorylation on multiple subunits of both 19S (Rpt3, Rpt5, Rpt6, Rpn6 and Rpn10) and 20S (α5 and α7) proteasome particles. Several studies have shown that phosphorylation of the proteasome subunits can either inhibit or stimulate the proteasome’s degradation activity on cell cycle related proteins, thereby affecting cell proliferation^6–9^. Given the complexity of eukaryotic cell cycle regulation network^10–12^, the detailed cell cycle regulation mechanism and its origin remain a mystery.

TACK superphylum in archaea is phylogenetically close to eukaryotes^13–15^. The archaea of the order Sulfolobales within the Thermoproteota (originally Crenarchaeota) phylum exhibit a eukaryotic-like cell cycle which contains G1, S, G2, M, and D (cell division) phases, accounting for < 5%, approximately 30%, > 50%, < 5%, and < 5% of the whole cell cycle, respectively. The cell division machinery in Sulfolobales resembles that of eukaryotes, encompassing several ESCRT-III (endosomal sorting complex required for transport III) homologs (also called CdvBs), Vps4 ATPase (or CdvC), and an archaeal-specific protein CdvA^16–19^. As other archaea, the information processing machineries of Sulfolobales are similar to those in eukaryotes^20–25^. Moreover, studies have revealed that Sulfolobales harbor a high-order level of chromosome organization in the form of A and B compartments, indicating a prototypical chromosome architecture of eukaryotes^26,27^. Therefore, Sulfolobales are excellent models for investigating the mechanisms and origins of eukaryotic-like cell cycle processes and their regulation. Investigation in *Sulfolobus acidocaldarius* revealed that the proteasome initiates cell cytokinesis by degrading the cell division protein CdvB during the cell cycle^28^. We and others have shown that the cell division in *Saccharolobus islandicus* is dictated by a transcription factor called aCcr1^29,30^. The protein binds to the promoters of dozens of genes including *cdvA* and represses their transcription. However, many aspects of these processes remain largely unexplored.

The proteasome is highly conserved in archaea^31^. Sulfolobales genomes encode one α subunit and two β subunits that make up the 20S core particle. The α subunits assemble into homo-heptameric rings whereas the two types of β subunits form hetero-heptameric rings^32^. In agreement with its role in cell division, the proteasome is crucial for the cells since all the subunit genes are likely essential in *Sa. islandicus*^33^. Homologs of proteasome assembly factors are present in archaea, some of which have been characterized^34,35^. However, it remains elusive how the proteasome is *do novo* assembled in archaeal cells and if this process is associated with phosphorylation-mediated regulation.

Archaeal protein kinases and phosphatase are more similar to their eukaryotic counterparts than to those found in bacteria^36^. Eukaryotic protein kinases (ePKs) can be classified into typical and atypical categories, with the former harboring 12 conserved classical subdomains whereas the later lacking sequence similarity^37,38^. The genomes of Sulfolobales encode 9–11 ePKs^36^. Studies have shown that archaeal atypical ePKs, such as Bud32, Rio1 and Rio2 homologs, perform conserved roles in tRNA modification and ribosome biogenesis, respectively, similar to their counterparts in eukaryotes^39,40^. In addition, it was reported that typical ePKs in Sulfolobales are involved in a complex regulation network that governs various cellular processes including cell mobility, biofilm formation, DNA damage response, and DNA repair^41–45^. However, it remains unclear whether archaeal ePKs play a role in cell cycle regulation.

In this study, we reveal that ePK2 in *Sa. islandicus* exhibits a cyclic expression pattern with a peak during cell division at the protein level. Additionally, the α subunit of the proteasome (PsmA) was identified as a substrate of ePK2. Phosphorylation of PsmA affects proteasome assembly and regulates its activity, thereby controlling cell division progression. This study has identified an innovative and presumably effective mechanism for regulating cell division in Sulfolobales archaea. This mechanism involves ePK-mediated proteasome phosphorylation and may be regarded as a prototype for the proteosome phosphorylation-mediated regulation of the eukaryotic cell cycle.

## Results

### The ePK2 gene is expressed cyclically in *Sa. islandicus*

In eukaryotes, protein phosphorylation plays a crucial role in cell cycle control^3^. Previous studies have shown that two ePKs, Saci_1193 and Saci_1694, in *S. acidocaldarius* were transcribed in a cell cycle-dependent manner^46^. Saci_1193 was significantly induced alongside genes related to chromosome segregation and cell division^46^. The genome of *Sa. islandicus* REY15A encodes a homolog of Saci_1193 (SiRe_2030). Transcriptomic analysis indicates that the transcription of *sire_2030* also changes in a cell cycle-dependent manner alongside with genes involved in chromosome segregation and cell division, while the transcription of other protein kinases does not exhibit a clear cyclical pattern^29^ (Fig. 1A and Fig. S1). To validate the cyclic pattern of ePK2, cells were synchronized at the G2 phase by treatment with acetic acid for 6 h. After the acetic acid was washed out, the cells were cultured in fresh medium and allowed to progreed through the cell cycle (Fig. 1B and 1C). Cell samples were collected at specified time points for RT-qPCR (Fig. 1D). The result revealed that *ePK2* mRNA gradually increases after release, peaking at about 2 h, consistent with the transcriptomic data. To assess whether the expression changes cyclically at protein level, Western blotting analysis was conducted and the results revealed that the SiRe_2030 level follows a cyclic pattern, peaking around 3 h after release (Fig. 1E). These results suggest that SiRe_2030, named as SisePK2 hereafter, and its homologs in Sulfolobales likely play a role in regulating cell cycle progression.

**Figure 1.**
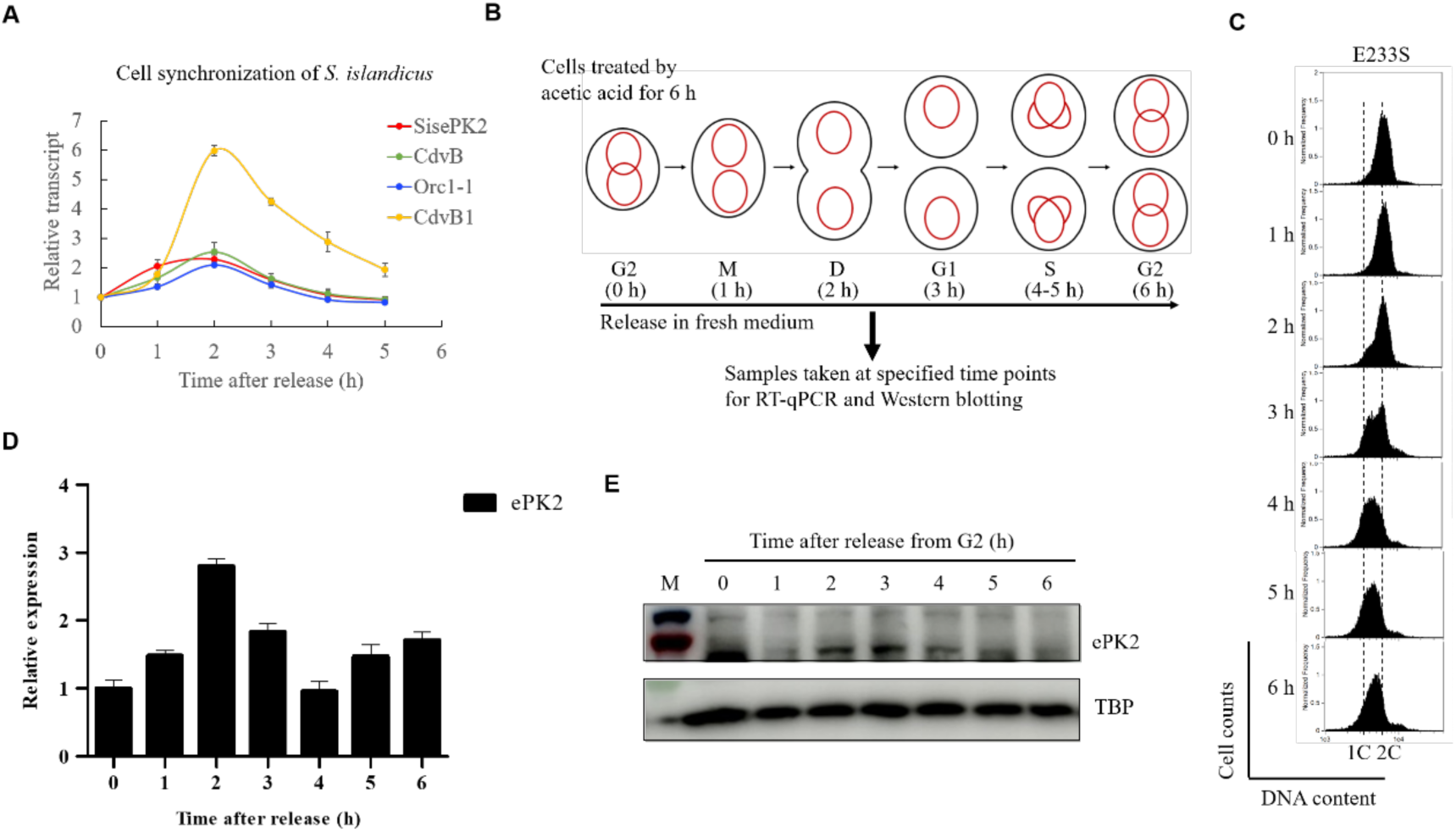
ePK2 exhibits cyclical expression. (**A**) Transcriptional pattern of ePK2 in the cell released from G2 phase. Data obtained from the transcriptomic analysis on synchronized samples of *Sa. islandicus* E233S (Yang *et al*., *Nucleic Acids Res*, 2023) show relative transcription levels of genes encoding for ePK2, DNA replication initiation factor Orc1-1, and two cell division proteins CdvB and CdvB1. Relative transcripts were calculated as FPKM (Fragments Per Kilobase transcript per Million mapped reads) of each gene divided by the level at 0 h, which was set as 1. Error bars represent standard deviations. (**B**) Schematic of the experimental workflow for cell cycle synchronization and time-point sampling of E233S cells. Cells were synchronized with acetic acid treatment for 6 h and released by washing with 20 mM sucrose and culturing in fresh preheated medium. Cells were collected at specified time points after release, and total RNA was isolated for RT-qPCR and the cell pellets were used for Western blotting. (**C**) Flow cytometry profiles of *Sa. islandicus* E233S after synchronization. At least 20,000 cells per sample were collected for the analysis. Dashed lines denote cells containing one and two copies of chromosome, 1C and 2C, respectively. (**D**) RT-qPCR verification of the cyclical transcriptional pattern of *ePK2* after synchronization. Data were normalized to 16S rRNA levels. Error bars represent standard derivations of three replicates. (**E**) Western blotting to detect ePK2 in the cells released from G2 phase. Samples at specified time points were analyzed by SDS-PAGE followed by Western blotting with protein-specific antibodies. TBP was used as a reference.

### Deletion of e*PK2* leads to impaired cell cycle progression

To investigate the role of ePK2 in cell cycle regulation, we constructed a deletion strain Δ*ePK2* (Fig. S2A and S2B) and characterized its phenotype by measuring cell growth and microscopy. Interestingly, we found that while asynchronized cells of Δ*ePK2* showed very slight difference in growth, cell morphology and cell cycle progression (Fig. 2A–2D), synchronized cells displayed distinct flow cytometry profiles compared with the wild type E233S (Fig. 2E). In the wild type cells, the number of G1 cells gradually increased at 2 h after release from G2 and peaked at 4 h. However, in Δ*ePK2* cells, the G1 peak was barely detected within 6 h after release, suggesting a deficiency in cell cycle progression caused by a failure in chromosome segregation or cell division, or a decoupling of cell division and the initiation of DNA replication. And in the strain Δ*ePK2*/pSe-NP-ePK2, G1 peak was recovered, confirming that the phenotype is strictly due to the deletion of *ePK2* (Fig. S3). In Δ*ePK2*/pSe-NP-ePK2, the *ePK2* deletion was complemented using the shuttle vector pSe-NP-ePK2, which was modified from pSeSD^47^, with the arabinose promoter being replaced with the native promoter (NP) of *ePK2*. To further investigate whether the absence of the G1 peak depends on the ePK2 kinase activity, we constructed a strain with in-frame catalytic deficient mutations, ePK2-D498A (Fig. S2C). The growth and cell morphology of asynchronized ePK2-D498A cells were not different from those of E233S and Δ*ePK2* (Fig. 2A–2D), but the flow cytometry profiles of the synchronized catalytic deficient strain resembled those of Δ*ePK2*, suggesting that the deficiency in cell cycle progression was due to the absence of ePK2 kinase activity (Fig. 2E). These results indicate that ePK2 and its kinase activity play an essential role in facilitating cell progression from G2 to G1 and suggest that phosphorylation of ePK2 target(s) may be involved in regulating chromosome segregation or cell division.

**Figure 2.**
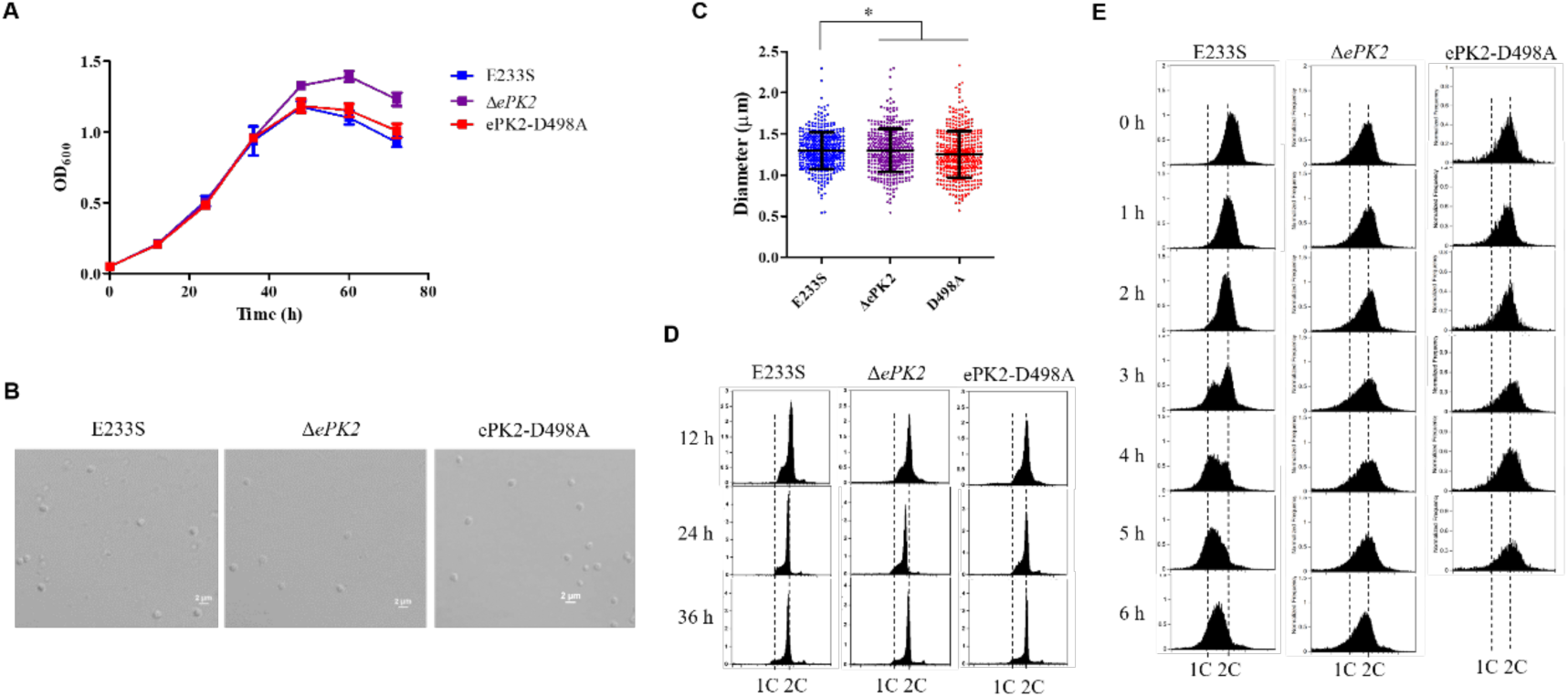
Elimination of *ePK2* and its catalytic activity impair cell cycle progression of synchronized cells. (**A**) Growth curves of Δ*ePK2* and ePK2-D498A. Cells were cultured aerobically at 75°C to OD_600_=0.4-0.8, and then transferred to fresh medium with an initial estimated OD_600_ of 0.05 for follow-up growth curve determination. (**B**) Microscopic analysis of Δ*ePK2* and ePK2-D498A. Samples were taken at early log phase (OD_600_∼0.3) and observed under a NIKON TI-E inverted fluorescence microscope. Scale bar, 2 μm. (**C**) Statistics of cell sizes in (B). Counted number (n) and average diameter (d) for each sample are as follows: E233S (n=398, d=1.296 ± 0.226), Δ*ePK2* (n=407, d=1.299 ± 0.260), and ePK2-D498A (n=402, d=1.253 ± 0.283). The values (mean ± SD) were obtained based on three independent biological repeats. Significance (*P* value) was calculated by the Wilcoxon test (two-tailed) using GraphPad. *n.s.*, not significant. (**D**) Flow cytometry analysis of *ΔePK2* and ePK2-D498A cells grown at 12 h, 24 h, and 36 h in (A). Samples were fixed by 70% ethanol overnight, washed with PBS buffer and stained with SuperGreen for DNA content analysis. At least 20, 000 cells were analyzed for each sample with over 90% cells from each sample included in the analysis. Dashed lines denote cells containing one and two copies of chromosome, 1C and 2 C, respectively. (**E**) Flow cytometry analysis of *ΔePK2,* and ePK2-D498A after synchronization by acetic acid treatment. Samples were taken at specified time points after released from G2 phase, fixed by 70 % ethanol and stained with propidium iodide (PI) for the analysis.

### Overexpression of ePK2 impairs cell division

To further explore the *in vivo* function of ePK2, we constructed an ePK2 overexpression strain based on the shuttle vector pSeSD with an arabinose-inducible promoter^47^. We found that overexpression of ePK2 inhibited cell growth and led to the formation of enlarged cells (2.332 ± 0.454 μm vs 1.241 ± 0.244 μm) (Fig. 3A–3C). Flow cytometry analysis showed that cells with multiple copies of chromosomes (>2C) were produced, indicating that although cell division was impaired, genome replication remain unaffected (Fig. 3D). To determine whether the inhibition was dependent on the kinase activity of ePK2, we constructed a strain overexpressing the point mutant of ePK2 on the conserved catalytic residue D498 (ePK2-D498A). Our results showed that the ePK2-D498A overexpression strain exhibited partially slower growth compared to the control harboring the empty vector but performed better than the ePK2 overexpression strain (Fig. 3A). Moreover, the average cell size and the ratio of >2C cells increased to a less extent compared to those of the ePK2 overexpression strain (1.241 ± 0.244 μm vs 1.941 ± 0.457 μm) (Fig. 3B–3D). These results suggest that the growth inhibition observed in the ePK2 overexpression strain was partially dependent on its kinase activity. Furthermore, our study revealed that the G1 peak of synchronized ePK2 overexpression strain was absent and the DNA content gradually increased after cells were released from G2 phase, while the G1 peak of synchronized ePK2-D498A overexpression strain gradually increased but in a slower speed than that of the strain with the empty vector (Fig. 3E). This indicates that overexpression of the wild type ePK2 significantly hinders chromosome segregation or cell division, but not genome replication. These results also imply that phosphorylation of ePK2 target(s) needs to be strictly regulated for normal cell cycle progression, such as accurate chromosome segregation/cell division or DNA replication initiation.

**Figure 3.**
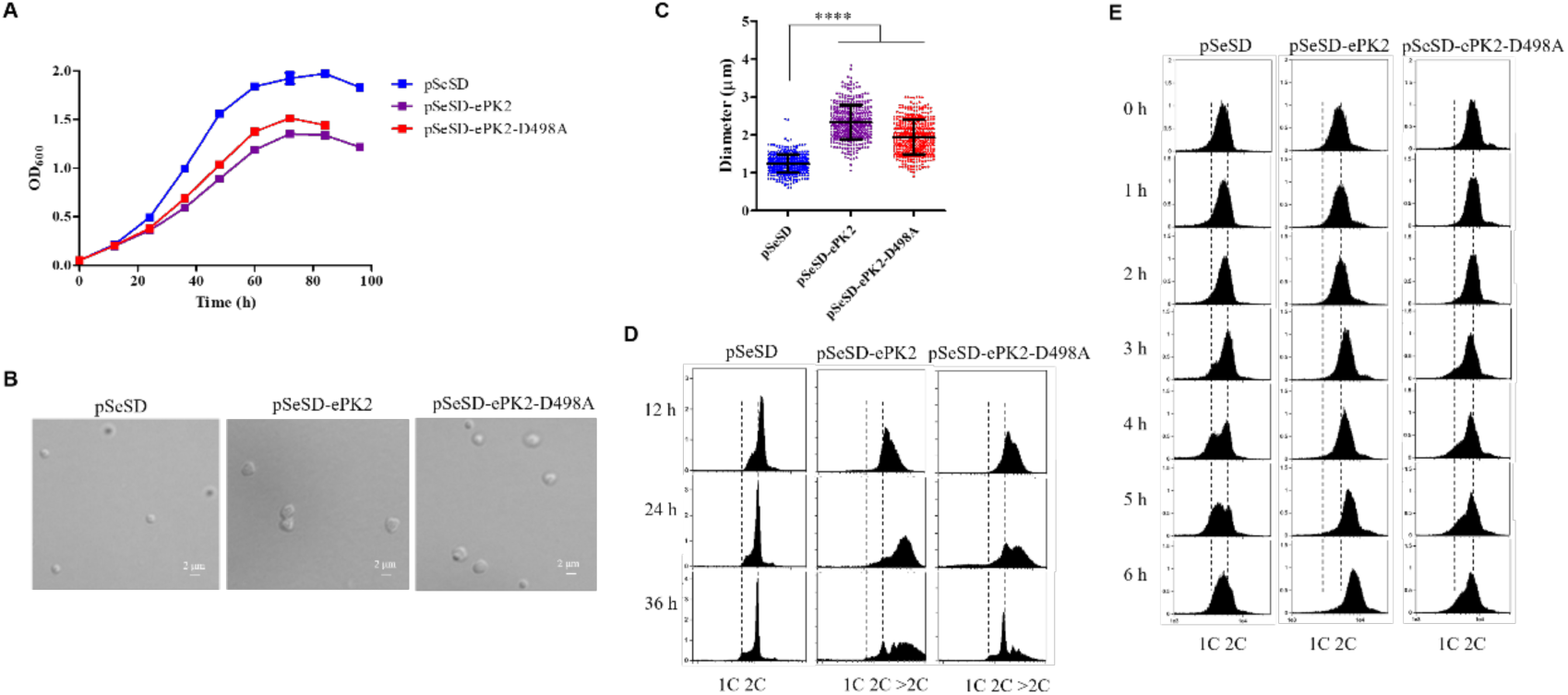
Overexpression of ePK2 hinders cell growth and cell division, resulting in the formation of enlarged cells. (**A**) Growth curves of strains overexpressing ePK2 and its kinase-dead mutant D498A. Protein induction was carried out in ATV medium using pSeSD-based vectors. E233S with the empty vector pSeSD was used as a control. (**B**) Microscopic analysis of strains overexpressing ePK2 and ePK2-D498A. Samples were taken at early log phase for microscopy. Scale bar, 2 μm. (**C**) Statistical analysis of cell sizes in (B). Cells carrying pSeSD (n=423, d=1.241 ± 0.244 μm), pSeSD-ePK2 (n=429, d=2.332 ± 0.454 μm) and pSeSD-ePK2-D498A (n=427, d=1.941 ± 0.457 μm) were analyzed. The values represent the mean ± SD from three independent biological repeats. Significance (*P* value) was calculated using the Wilcoxon test (two-tailed) in GraphPad. ****, *p*<0.0001. (**D**) Flow cytometry of strains overexpressing ePK2 and ePK2-D498A. Samples were taken at 12, 24, and 36 h after being cultured in ATV medium. (**E**) Flow cytometry of the ePK2 and ePK2-D498A overexpression strains after release from G2 phase. Cells were synchronized with acetic acid treatment for 6 h, followed by washing out the acetic acid and culturing the cells in ATV medium. The samples were taken at specified time points for flow cytometry analysis with at least 20, 000 cells per sample.

### The α subunit (PsmA) of *Sa. islandicus* proteasome is a substrate of ePK2

To identify potential substrates or interactors of ePK2, His-tagged ePK2 was expressed in *Sa. islandicus* using the overexpression strain and purified by Ni-NTA and gel filtration. SDS-PAGE analysis of the samples identified four co-purified bands in fraction 11, preceding the ePK2 peak (fraction 12) (Fig. 4A). Mass spectrometry (MS) analysis revealed that the co-purified proteins included proteasome α subunit (PsmA), thermosomes (SiRe_1214 and/or SiRe_1716), and fructose-1,6-biphosphate aldolase/phosphatase (Fbp). To determine whether ePK2 directly phosphorylates PsmA, an *in vitro* kinase assay was performed using purified proteins from *E. coli* (Fig. 4B and S4). Phosphorylated proteins were detected by Western blotting with an antibody specific for phosphorylated Thr/Tyr. The result showed that the wild type ePK2, but not the kinase-dead mutant ePK2-D498A, was able to phosphorylate PsmA (Fig. 4B). Additionally, the intensities of phosphorylation signals increased with increasing concentrations of PsmA. Furthermore, we also detected the PsmA phosphorylation by another protein kinase ePK1 (SiRe_2056), known for its higher activity with a broad substrate range *in vitro*^45^. As illustrated in Fig. 4C, the phosphorylation signal of PsmA by ePK1ΔTM (ePK1 lacking its transmembrane domain) was comparable to that observed with ePK2. These results indicate that, *in vivo*, PsmA was probably mainly phosphorylated by ePK2 but not ePK1, which would be anchored on the cellular membrane.

**Figure 4.**
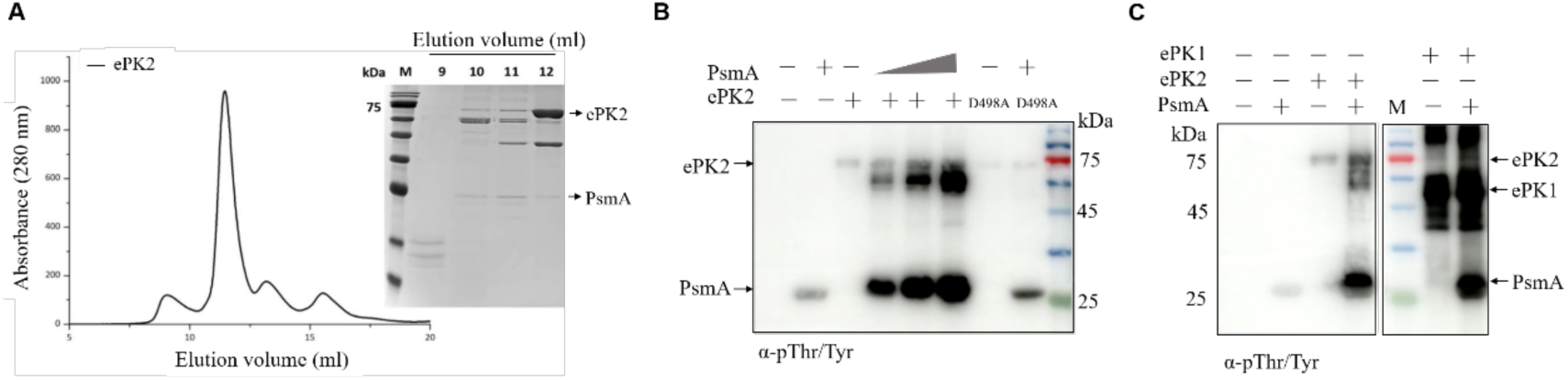
The α subunit of proteasome, PsmA, is a substrate of ePK2. (**A**) Purification of ePK2 from its overexpression strain. Cells were cultured in STV medium for low induction and His-tagged ePK2 was purified with Ni-NTA and gel filtration. The elution profile and SDS-PAGE analysis are shown with bands of ePK2 and PsmA indicated. (**B**) *In vitro* phosphorylation of PsmA by ePK2 and its kinase-dead mutant (D498A). Different concentrations of PsmA (0.5, 1 and 2 μM) were incubated with 0.5 μM ePK2 or ePK2-D498A (with 2 μm PsmA) at 65°C for 30 min and analyzed by Western blotting using the phosphorylated Thr/Tyr antibody. (**C**) ePK1ΔTM (ePK1 lacking its transmembrane domain) phosphorylates PsmA at a similar activity compared to ePK2. ePK1 and ePK2 was individually mixed with 2 μM PsmA and incubated at 65°C for 30 min. The PVDF membranes were extracted from the same membrane.

### Overexpression of ePKs results in accumulation of cell division proteins

Previous studies have shown that cyclically expressed cell division protein CdvB is a target of the proteasome, and failure to degrade CdvB by proteasome impairs cell division in both *S. acidocaldarius* and *Sa. islandicus*^19,28^. This suggests that proteasome activity is crucial for cell division in Sulfolobales. Since PsmA is a substrate of ePKs, we hypothesized that overexpression of ePK2 might affect the proteasome activity, leading to deficiency in cell division. To test this, we compared the CdvB levels in synchronized cells of wild type and ePK2 overexpression strains using Western blotting (Fig. 5A). In wild type cells, CdvB accumulated at 2–3 h after release from G2 and was rapidly degraded after 3 h, coinciding with completion of cell division in most cells. In contrast, in cells of the overexpression strain, the degradation of CdvB was apparently delayed until 5 h after release from G2, indicating that the proteasome activity was impaired in this strain. In addition, we also examined the levels of other cell division protein, CdvB1 and CdvB2, which work in conjugation with CdvB^18,19,28^. Treatment with Bortezomib, a proteasome-specific inhibitor, resulted in accumulation of CdvB1 and CdvB2, indicating they are also targets of the proteasome (Fig. S5A). Notably, we found that although CdvB1 displayed cyclic expression pattern, its degradation was delayed and slower compared to that of CdvB (Fig. 5A), consistent with its role at late stage in cell division^18,19,28^. As expected, degradation of CdvB1 was also inhibited upon ePK2 overexpression (Fig. 5A). Next, we detected the levels of CdvB1 and CdvB2 in asynchronized culture of ePK2 overexpression strain. As depicted in Fig. 5B, the overexpression strain exhibited approximately 8 times higher levels of ePK2 compared to the wild type strain and the levels of both CdvB1 and CdvB2 were significantly increased with ePK2 overexpression. To rule out the possibility that the accumulation of cell division proteins in the cell was a result of increased *cdvB1/cdvB2* transcription, we assessed both mRNA and protein levels at different time points following ePK2 induction. We observed that CdvB1 protein increased by approximately 2-fold after 6 h of induction, while its mRNA level only slightly increased after 8 h of induction and even decreased before 8 h (Fig. S6). These finding suggest that the elevation in CdvB1 protein level was attributed to its reduced degradation by the proteasome rather than increased transcription. In addition, the protein levels of two proteasome subunits, PsmA (SiRe_1271) and PsmB1 (SiRe_1237), were comparable to those of the wild type (Fig. 5B), suggesting that the proteasome content in the ePK2 overexpression strain remained unchanged.

**Figure 5.**
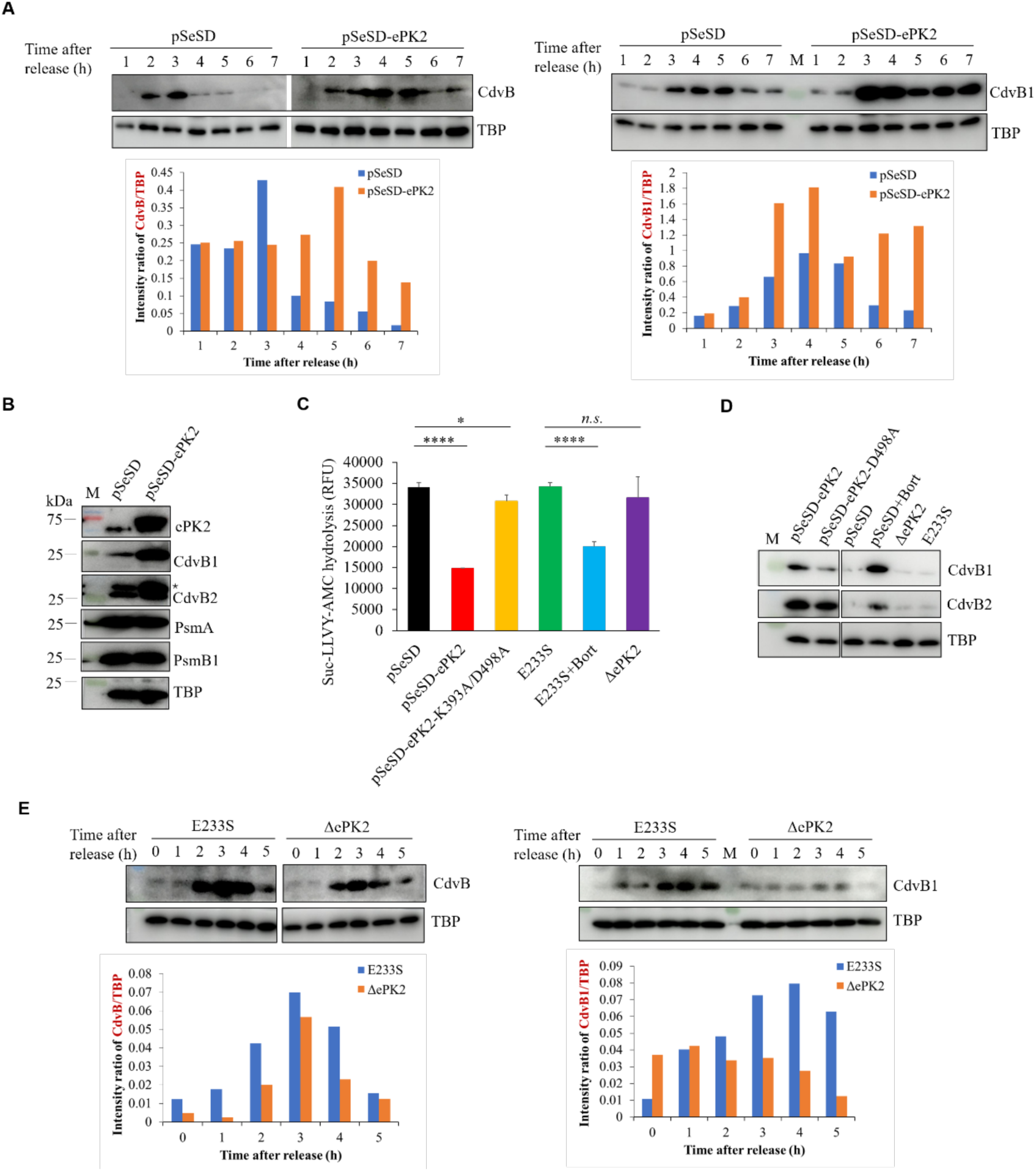
Overexpression of ePK2 inhibits proteasome activity while its deletion stimulates the activity *in vivo*. (**A**) Impaired degradation of CdvB and CdvB1 in synchronized ePK2 overexpression strain. Samples were taken after the cells were released from G2 phase and subjected to Western blot analysis using specific antibodies. Quantification of CdvB and CdvB1 using TBP as the control is shown at the bottom. (**B**) Accumulation of cell division proteins after ePK2 overexpression. Samples were taken at early log phase. The asterisk indicates the position of CdvB2. (**C**) Decreased *in vivo* proteasome activity in asynchronized cells with ePK2 overexpression but not *ePK2* deletion. Proteasome activities of whole cell lysate (WCL) of ePK2 overexpression and deletion strains were measured using the fluorescent oligopeptide Suc-LLVY-AMC as the substrate. The reaction mixture was incubated at 70°C for 30 min and detected by microplate spectrometer. E233S WCL treated with the proteasome specific inhibitor Bortezomib was served as a positive control. Significance (*P* value) was determined using a one-tailed *t* test in GraphPad. ****, *p*<0.0001; *n.s.*, no significant. (**D**) Inhibition of CdvB1 and CdvB2 degradation of cells with ePK2 overexpression but not *ePK2* deletion. pSeSD based vectors were used for protein overexpression, with ePK2-D498A and ePK2-K393A/D498A being catalytic dead mutants. (**E**) Enhancement of CdvB and CdvB1 degradation in synchronized *ΔePK2* cells. Quantification of CdvB and CdvB1 using TBP as the control is shown at the bottom. The separated blot with the same antibody in (D) and (E) were extracted from same membranes.

### ePK2 phosphorylation inhibits cellular proteasome activity and Δ*ePK2* exhibits accelerated degradation of the cell division proteins

To determine whether the proteasome activity was impaired in the ePK2 overexpression strain, we measured the proteasome activity of whole cell lysate using a fluorescent substrate, Suc-LLVY-AMC ^48^. Our finding revealed that the proteasome activity of the total soluble cell extract was significantly reduced in the ePK2 overexpression cells as compared with that in cells harboring the empty vector pSeSD (Fig. 5C). In addition, the proteasome activity of ePK2-K393A/D498A overexpression strain was only slightly weaker than the control, consistent with our previous result that the inhibition was partially dependent on the kinase activity of ePK2 (Fig. 5C). Furthermore, Western blotting showed that both CdvB1 and CdvB2 accumulated in ePK2 overexpression strain, while the ePK2-D498A overexpression only led to partial accumulation of these proteins (Fig. 5D). As expected, in Δ*ePK2* neither the proteasome activity nor accumulation of CdvB1 and CdvB2 was significantly different from those in strain E233S, in agreement with the result that the phenotype of ΔePK2 cells was similar to that of E233S under asynchronization conditions (Fig. 5C and 5D). This observation raises the possibility that there might be other protein kinase(s), such as ePK1, playing redundant role in the asynchronized cells. Taken together, we conclude that overexpression of ePK2 inhibits cellular proteasome activity in *Sa. islandicus*.

To further investigate the impact of the absence of ePK2 on cell division, we examined levels of cell division proteins in synchronized Δ*ePK2* cells releasing from G2 phase using Western blotting. Strikingly, levels of CdvB and CdvB1 were significantly reduced in synchronized Δ*ePK2* cells compared to the wild type E233S (Fig. 5E). Moreover, cells with a catalytic deficient mutation, ePK2-D498A, also exhibited lower levels of CdvB1 than E233S after synchronization (Fig. S5B). These results indicate that, opposite to ePK2 overexpression, cells without ePK2 may lead to abnormally high proteasome activity resulting in failure in accumulation of CdvB proteins post-G2 phase and subsequent impaired cell division (Fig. 2E).

### ePK2 phosphorylates PsmA at S200 and T213

To identify the phosphorylated residues of PsmA by ePK2, we performed comparative quantitative phosphoproteomic analysis between ePK2 overexpression strain and the control. The result showed that phosphorylation of 434 sites (from 305 proteins) increased by over 2-fold in ePK2 overexpression strain, among which 43 residues (from 38 proteins) increased by over 10-fold (Supplementary dataset). Thermosomes, Fbp, and PsmA are proteins with phosphorylation sites increased over 10-fold. Four residues (S46, S59, S200, and T213) on PsmA were phosphorylated and the phosphorylation at S200 and T213 increased by 2.7-fold and 11.7-fold, respectively, with ePK2 overexpression (Fig. 6A, Table 1). To verify these residues were phosphorylated, wild type PsmA and its single, double or multiple point mutants were purified from *E. coli* (Fig. S4) and subjected to a kinase assay with ePK2. Radiolabeled [γ-^32^P] ATP was used to quantify the phosphorylation levels. As shown in Fig. 6B and 6C, although phosphorylation of S200A (102.0%) did not decrease compared to the wild type PsmA, phosphorylation of T213A was reduced to 75.7% and that of the double mutant S200A/T213A decreased further to 38.8%, suggesting that ePK2 phosphorylated both S200 and T213 of PsmA. However, phosphorylation of S46A/S59A/S64A/T213A (78.1%) and S46A/S59A/S64A/S200A/T213A (51.0%) did not decrease further, compared to those of T213A and S200A/T213A, respectively, indicating that S46A/S59A/S64A is not the targets of ePK2 *in vitro*. Alternatively, there could be interplay between phosphorylated sites. Taken together, we revealed that ePK2 phosphorylates PsmA at both S200 and T213 *in vivo* and *in vitro*.

**Figure 6.**
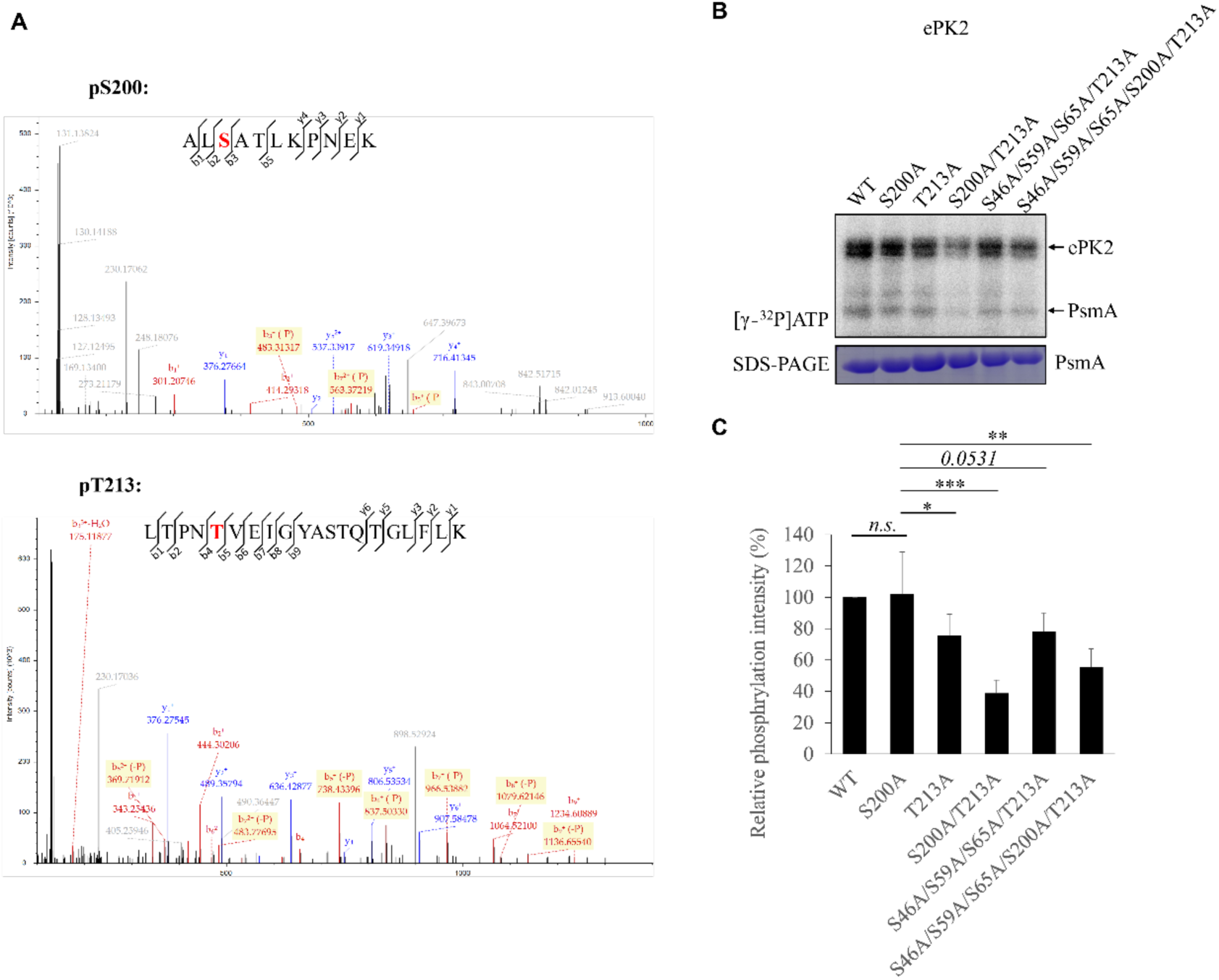
ePK2 phosphorylates PsmA at S200 and T213. (**A**) Mass spectra of two oligopeptides containing phosphorylated S200 and T213. The phosphorylated residues are indicated in red. (**B**) ePK2 phosphorylation on PsmA and its point mutants. The *in vitro* kinase reaction was performed as that in Figure 4B except that [γ-^32^P]ATP was used instead of cold ATP. Upper panel, radiolabeled exposure of [γ-^32^P]ATP signal; low panel, SDS-PAGE of PsmA and its mutants. A representative gel is shown. (**C**) Quantitative results of (B). The data show relative phosphorylated ratios between the mutant and wild type (WT) PsmA proteins based on five independent experiments. Significance (*P* value) was calculated using a one-tailed *t* test in GraphPad. *, *p*<0.05; **, *p*<0.01; ***, *p*<0.001; *n.s.*, not significant.

**Table 1.** Identification of phosphorylated residues in PsmA in the phosphorproteome analysis of ePK2 overexpression strain.

| **Position of PsmA** | **Motif <sup>a</sup>** | **pSeSD** | **pSeSD-ePK2** | **Fold Change** |
| --- | --- | --- | --- | --- |
| T213 | EKLTPN**T**VEIGYA | 149271.2 | 1743609.1 | 11.68081 |
| S200 | LALKAL**S**ATLKPN | 525092.97 | 1404456.6 | 2.674682 |
| S59 | EKRKAQ**S**LLDVDS | 582475.81 | 555168.13 | 0.953118 |
| S46 | IGIKSK**S**GVVIAS | 884005.76 | 415292.04 | 0.469784 |

<sup>a</sup>, the phosphorylated sites are indicated in red.

### Overexpression of PsmA phospho-mimic mutants S200D and T213E has pronounced and modest impacts on cell division, respectively

To explore the functional mechanism of PsmA phosphorylation, we initially attempted to delete *psmA* in *Sa. islandicus* E233S. Consistent with previous findings from a genome-wide transposon mutagenetic analysis^33^, we were unable to obtain a *psmA* knockout strain, confirming its essential role in cell viability. Next, we constructed strains overexpressing the wild type PsmA and its phosphorylation site mutants including dephosphorylation mutants S200A, T213A, and S200A/T213A, as well as phospho-mimic mutants S200D, T213E, and S200D/T213E. Increased levels of PsmA proteins (approximately 3–5 folds) were observed in the overexpression strains compared to cells harboring the empty vector (Fig. S7). As depicted in Fig. 7A and 7B, overexpression of wild type PsmA resulted in growth retardation and cell enlargement (1.560 ± 0.287 μm vs 1.274 ± 0.155 μm), although to a lesser extent compared to the ePK2 overexpression strain. In addition, PsmA overexpression did not affect genome replication since the enlarged cells were found to contain more than 2 copies of chromosomes as revealed by flow cytometry (Fig. 7C). These results indicate that PsmA overexpression impaired the proteasome activity and inhibited cell division, reminiscent of the phenotype of the ePK2 overexpression strain. To validate this, we examined the CdvB1 level and proteasome activity in the PsmA overexpression cells. As anticipated, the CdvB1 level increased and the proteasome activity was inhibited (Fig. 7D and 7E). Furthermore, CdvB1 degradation was significantly slower in the PsmA overexpression strain compared to the wild type, as revealed by Western blotting of synchronized cells (Fig. S8), suggesting a deficiency in proteasome-mediated degradation. We assume that PsmA overexpression may affect correct proteasome assembly and impair its activity *in vivo* (see below).

**Figure 7.**
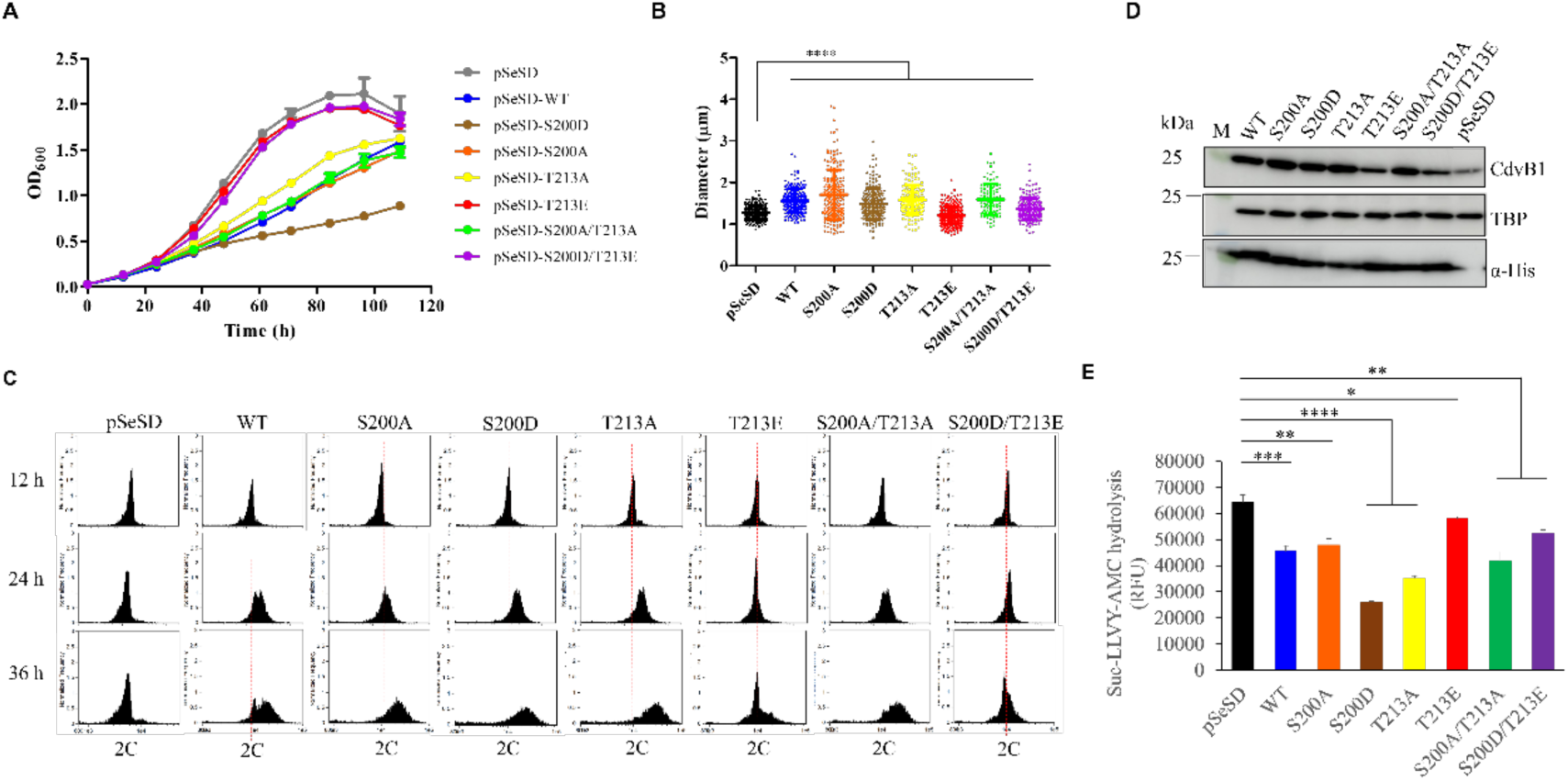
Overexpression of PsmA phospho-mimic mutants S200D and T213E confers different effects on the cell. (**A**) Growth curves of PsmA and its mutant overexpression strains. Cells were cultured in STV medium for 3 times before inoculation in ATV medium to OD_600_∼0.03 for protein induction. (**B**) Statistical analysis of cell sizes of PsmA and its mutant overexpression strains. Cell number counted (n) and average diameter (d) for each strain are as follows: pSeSD (n=280, d=1.274 ± 0.155 μm), WT (n=198, d=1.560 ± 0.287 μm), S200A (n=206, d=1.706 ± 0.599 μm), S200D (n=197, d=1.490 ± 0.370 μm), T213A (n=172, d=1.587 ± 0.356 μm), T213E (n=315, d=1.211 ± 0.206 μm), S200A/T213A (n=97, d=1.596 ± 0.376 μm) and S200D/T213E (n=196, d=1.366 ± 0.258 μm). The values represent the mean ± SD from three independent biological repeats. Significance (*P* value) was calculated using the Wilcoxon test (two-tailed) in GraphPad. ****, *p*<0.0001. (**C**) Flow cytometry profiles of PsmA and its mutant overexpression strains. Cells were cultured in ATV medium and taken at specified time points for flow cytometry. At least 20, 000 cells per sample were analyzed. (**D**) Detection of CdvB1 levels in the overexpression strains of PsmA and its mutants. CdvB1 and PsmA proteins were detected by CdvB1 antibody and a His-tag antibody, respectively, with TBP as a control. (**E**) The proteasome activities of PsmA and its mutant overexpression strains. Significance (*P* value) was determined by the *t* test (one-tailed) using GraphPad. *, *p*<0.05; **, *p*<0.01; ***, *p*<0.001; ****, *p*<0.0001.

Surprisingly, all dephosphorylated mutant overexpression cells (S200A, T213A, and S200A/T213A) exhibited a similar phenotype to that of wild type overexpression cells (Fig. 7). Intriguingly, overexpression of phospho-mimic mutants, S200D and T213E, had contrasting effects on the cells: S200D overexpression further inhibited cell growth and cellular proteasome activity, while T213E overexpression basically did not influence the growth and CdvB1 degradation, and its proteasome activity was only slightly reduced as compared with that of Sis/pSeSD (Fig. 7 and Fig. S6). Moreover, overexpression of the double phospho-mimic mutant S200D/T213E had the same effect as T213E (Fig. 7 and Fig. S8), indicating that phospho-mimic mutation of T213 somehow neutralized the severe effect of S200D on the cell. We speculate that mutation of S200 and T213 to alanine did not affect their inter-subunit interaction and interactions with other factors, resulting in a phenotype similar to that of the WT overexpression strain. On the other hand, mutation of S200 and T213 to negative-charged phospho-mimic residues abolished all or part of these interactions. In addition, the effect of T213E may be upstream of that of S200D.

### Phosphorylation of residues S200 and T213 does not affect the interaction between α subunits

In order to elucidate the different effects of S200D and T213E overexpression on cells and uncover potential regulatory mechanism through PsmA phosphorylation, we isolated proteasomes in cells overexpressing His-tagged wild type PsmA and its mutants, and characterized them by SDS-PAGE, Western blotting, and microscopy with a Focused Ion Beam Scanning Electron Microscope. Notably, we were able to observe the co-purified non-tagged PsmA chromatin gene-encoded band and the quantity of non-tagged PsmA co-purified with His-tagged T213E was apparent lower than that of the His-tagged PsmA (Fig. S9A). However, the migration of His-tagged S200D and non-tagged PsmA were too close to be differentiated. In addition, one of the two β subunits, PsmB1, were also detected in all three samples (Fig. S9B). Electron microscopy analysis revealed that the majority of proteasome particles consisted of two heptameric rings, instead of 20S core particle with four rings, despite the presence of the β subunit in these samples (Fig. S9C-9E). This was further confirmed by analyzing PsmA purified from *E. coli*, yielding the same result (Fig. S9F). This suggests that the two phosphor mimic mutations, or phosphorylation at these two residues, do not affect the interaction between α subunits.

### Phosphorylation at S200 abrogates 20S-PAN assembly whereas phosphorylation at T213 may affect *de novo* 20S assembly

To gain deeper insights into the functional mechanism of S200 and T213 phosphorylation, we conducted an investigation into the assembly of 20S and 26S (20S with proteasome-activating nucleotidase (PAN)) through sucrose density gradient centrifugation followed by proteasome activity analysis. The result of the Western blotting analysis revealed two signal peaks of PsmB1 in the strain carrying the empty vector, observed at fraction 2 and fractions 8–9, respectively (Fig. 8A). The proteasome activity assay showed that the fractions corresponding to these two signal peaks exhibited the highest peptidase activities, indicating that the fractions 2 and 8–9 contained 20S and 26S proteasomes, respectively (Fig. 8B). To confirm that the observed activity was not attributed to other peptidases in the cell, various protease inhibitors, including PSMF, protease inhibitor mixture, and a proteasome-specific inhibitor Bortezomib, were added in the assay. It was found that Bortezomib but not any other protease inhibitor suppressed the activity (Fig. 8B). Furthermore, addition of ePK2 in the reaction mixtures led to a partial inhibition of the protease activities in both fractions, consistent with that phosphorylation of 20S proteasome subunit impair its functionality (Fig. 8B).

**Figure 8.**
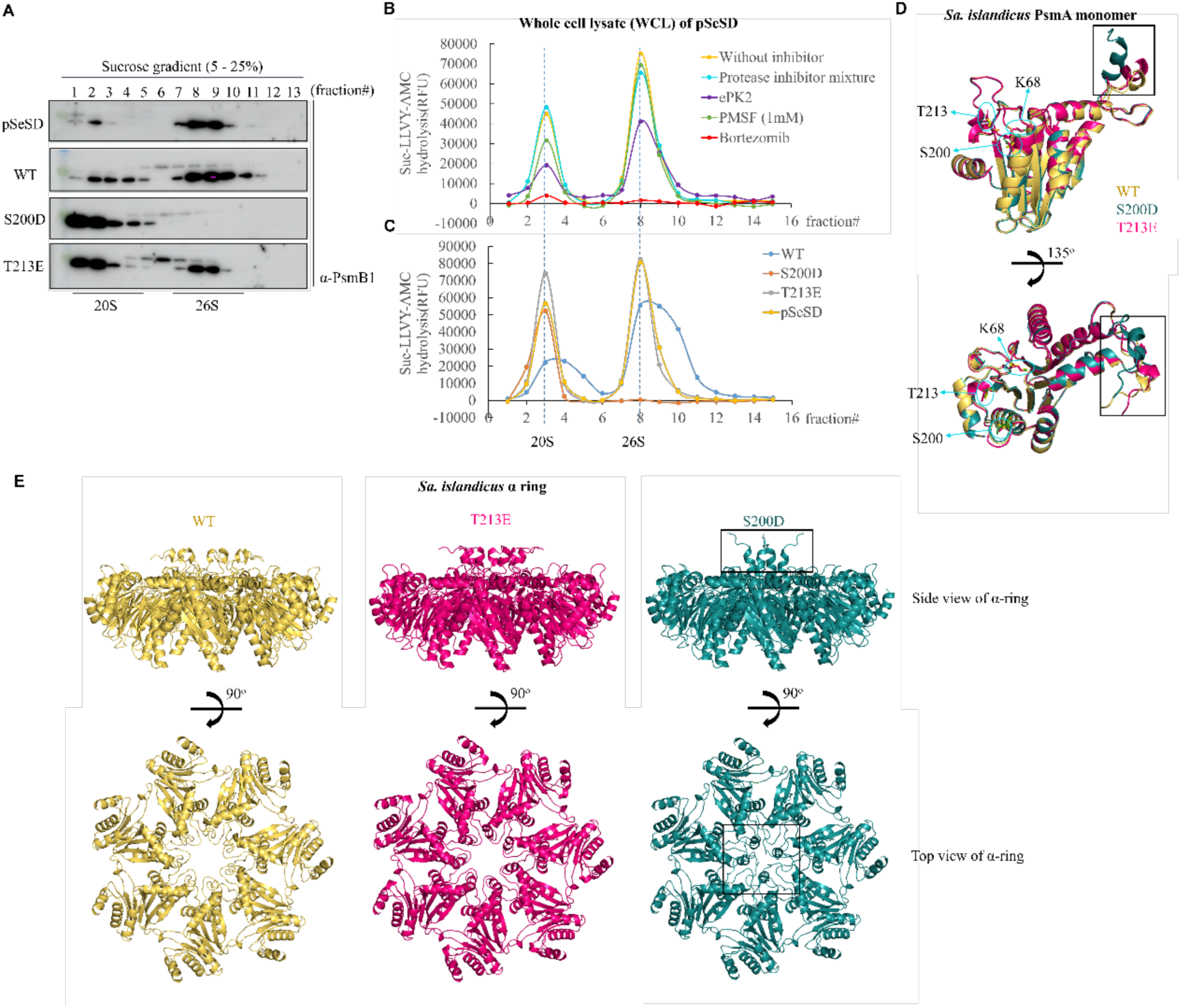
Overexpression of S200D eliminates 26S assembly, while overexpression of T213E has no effect on cellular 20S or 26S assembly. (**A**) The isolation of the 20S and 26S proteasomes from cells overexpressing PsmA. A 0.5 ml aliquot of soluble whole cell lysate (WCL) of PsmA overexpression strains was centrifuged with sucrose gradients ranging from 5%–25%. Samples were collected at 1 ml intervals and analyzed by Western blotting with PsmB1 antibody. The positions of 20S and 26S proteasomes are indicated at the bottom of the figure. (**B**) Proteasome activities of WCLs from cells carrying pSeSD and pSeSD-ePK2 with or without inhibitors. (**C**) Proteasome activities of WCLs from pSeSD-PsmA (WT), S200D and T213E. The positions of 20S and 26S proteasomes are indicated at the bottom of the curves. (**D**) Superposition of PsmA (WT) (yellow), S200D (cyan), and T213E (pink) monomers. The structures were predicted by AlphaFold 3. Phosphorylated residues, S200 and T213, and the conserved K68 interacting with HbYX motif are indicated by circles. The black rectangle highlights the conformational change in S200D. (**E**) Comparison of WT (yellow), S200D (cyan), and T213E (pink) heptamer rings. The upper panel shows a side view and the lower panel shows a top view, with black rectangles indicating conformational changes of S200D within the α ring.

We observed that the specific fractions containing 20S/26S proteasomes in wild type PsmA overexpression cells exhibited lower peptidase activities compared to cells carrying the empty plasmid (Fig. 8C). This suggests that overproduction of PsmA somehow disrupts the assembly of normal 20S and 26S proteasomes, resulting in the formation of 20S/26S proteasomes with decreased activities. This may explain the observed phenotype in the wild type PsmA overexpression strain. Strikingly, in S200D overexpression strain, only the 20S proteasome peak was present while the 26S proteasome peak was completely absent (Fig. 8A). The proteasome activity assay also showed a lack of peptidase activity at the position where the 26S proteasome typically appears (Fig. 8C), suggesting a failure of the 26S proteasome assembly in the cell. It has been demonstrated that the activation of 20S proteasome is triggered by the interaction of an HbYX motif at the C-terminus of PAN with the conserved Lys of PsmA, leading to the opening of 20S gate^49–51^. As a control, a strain overexpressing PsmA with a mutation at the conserved Lys, K68A, was constructed. As expected, we observed a significant decrease in both the 26S proteasome signal and activity peaks in K68A overexpression strain, similar to what was observed in the S200D overexpression strain (Fig. S10A and S10B). We also detected the sediments and peptidase activities of 20S/26S proteasomes in the ePK2 overexpression strain. The results showed that the protein peak and activity of the 26S proteasome were much lower than those in the control with the empty vector (Fig. S10C and S10D). This supports that elevated level of ePK2 with corresponding elevation of the proteasome phosphorylation disrupts the 26S proteasome assembly in the cell.

On the other hand, we found that the sediments and activities of the 20S and 26S proteasomes in the T213E overexpression strain were comparable to those observed in the Sis/pSeSD strain (Fig. 8A and 8C), aligning with their corresponding phenotypes (Fig. 8). The results imply that overproduced T213E in the cell could not be incorporated into the 20S/26S proteasomes. This was consistent with the result that purified T213E from its overexpression strain only pulled down lesser amounts of genome-encoded PsmA (Fig. S9A), suggesting that T213E formed α rings on its own but faced challenges in being integrated into functional internal α rings assembled *de novo*. Consistently, the analysis of S200D/T213E overexpression strain using sucrose density gradient centrifugation revealed a pattern identical to that of T213E and the control strain with the empty vector, indicating that failure of S200D/T213E to be incorporated into functional internal α rings due to the T213E mutation would erase the effect of S200D mutation (Fig. S10A and S10B). In summary, our findings suggest that phosphorylation of S200 abrogates 20S-PAN assembly, whereas phosphorylation of T213 may affect *de novo* α ring assembly.

### Structural modelling of the wild type and mutant PsmA proteins and the α rings

To investigate the potential molecular mechanisms behind the impact of S200 and T213 phosphorylation on the assembly of 20S/26S proteasome in *Sa. islandicus*, the structures of PsmA carrying these mutations and the assembled α rings were generated using Alphafold3. The monomeric structure T213E mutant closely resembled that of the wild type PsmA, with a slight misalignment at its N-terminus being the only difference (Fig. 8D). However, S200D exhibited a notable elongation of the α helix at the N-terminus with a further dislocation. In the predicted whole α rings, the wild type PsmA and T213E assembly are almost the same with the N-termini of seven monomers being arranged as a lid-like structure above the central channel of proteasome (Fig. 8E). On the other hand, the N-terminal α helix of S200D shifts deeper into the central channel, causing a disruption in the proper alignment of the N-termini from each monomer above the channel. This resulted in a structure with only four N-termini being visible while three are predicted to be buried down into the channel as loops (Fig. 8E). The structural change in the N-terminus of S200D may impede the interaction between PsmA and PAN, leading to a failure in the assembly of 26S proteasome. So far, the reason for the inability of T213E to be incorporated into α ring remain unclear, since there is minimal disparity observed in the structural models. This may suggest that there is a complex *de novo* assembly process of the α ring or the 20S proteasome which requires additional assembly factors *in vivo*. Supporting this notion, we have identified two potential 20S proteasome assembly factors (SiRe_1724 and SiRe_1163) that co-purified with PsmA in mass spectrometry analysis of *in vivo* Co-IP samples. Moreover, the interactions of PsmA with both assembly factors were confirmed by *in vivo* pull-down using overexpression strains (Fig. S11). The mechanisms behind the assembly of the 20S/26S proteasome in *Sa. islandicus* warrant further investigation in future studies.

## Discussion

Cells of the archaeal order Sulfolobales exhibit distinct cell cycle phases and serve as excellent models for investigation into cell cycle regulation mechanisms in early life forms. Previously, it has been shown that levels of cell division proteins in Sulfolobales are controlled by proteasome-mediated degradation and by the cyclic expression of the transcription factor aCcr1^28,29^. Here, we demonstrate that a cyclically expressed ePK-mediated phosphorylation plays an important role in regulating proteasome assembly. The cell cycle-dependent phosphorylation influences the degradation activity of the proteosome towards proteins like CdvBs and dictates cell cycle progression in *Sa. islandicus*. The finding of this study implies that Sulfolobales archaea have a prototype of phosphorylation-mediated cell division regulation mechanism.

Multiple phosphorylation residues in PsmA including S200 and T213 were identified through phosphoproteomic analysis of ePK2 overexpression cells. However, attempts to identify phosphorylated S200 and T213 of PsmA purified from asynchronized culture using Co-IP with anti-PsmA were unsuccessful, suggesting that this type of phosphorylation may be transient and limited to a specific period during the cell cycle (late M phase). Rapid dephosphorylation by active phosphatases likely follows to ensure precise control and efficiency in cell division^52^. This regulation is essential since the window for cytokinesis during cell division in Sulfolobales archaea is extremely short, e. g. only less than 7 min in *S. acidocaldarius*^53^.

In eukaryotes, the assembly of 20S proteasome is a complex process that involves several chaperons, Pba1–4 and Ump1 in yeast (PAC1–4 and POMP in mammals). The Pba1-Pba2 complex is responsible for appropriate incorporation of α subunits into α ring and prevents premature binding of the 19S particle to the α ring^54–56^. Additionally, Pba3-Pba4 and Ump1 are required for incorporation of β subunits and Ump1 prevents inappropriate dimerization of two half-proteasomes^57–59^. Recent Cryo-EM analysis has revealed the order of incorporation of α and β subunits during human 20S proteasome assembly. It was shown that PAC1-PAC4 have extensive contacts with α5, α6 and α7 with the C-termini of PAC1 and PAC2 inserting into the HbYX binding pockets between α subunits^60^. Our results suggest that phosphorylated T213 may be not incorporated into α ring during its *de novo* assembly. Indeed, a very limited amount of genome-encoded PsmA was pulled down when T213E was purified *in vivo* compared to WT (Fig. S9A). Although assembly of an archaeal 20S particle can be accomplished by co-expression of α and β subunits in *E. coli* without additional partners, the characterization of Pba homologs suggested that they may play a role in the assembling process in archaea^34,35^. Interestingly, two putative Pba1/2 homologs, SiRe_1163 and SiRe_1724, were identified in our MS analysis of the Co-IP samples using PsmA antibody and their interaction with PsmA was confirmed by *in vivo* pull-down assay (Fig. S11), suggesting that 20S proteasome assembly may also need chaperons in archaeal cells. Since Pba/PAC complexes interact with multiple α subunits through HbYX binding pockets in both archaea and eukaryotes and T213 locates at a loop adjacent to the HbYX binding pocket (Fig. 8D), mutation of T213E may disrupt the interaction of PsmA with the putative Pba complex and prevent the incorporation of T213E into the *de novo* α ring. Several lines of evidence from this study together strongly suggest that phosphorylation at S200 blocks the interaction between the 20S particle and PAN, resulting in a failure in 26S assembly (Fig. 7, Fig. 8 and Fig. S10), while phosphorylation of T213 may block α ring assembly. Further investigation is required to fully understand the role of these residues in 20S/26S assembly.

In eukaryotes, proteasome activities are regulated by phosphorylation of various subunits. For example, T25 of the 19S subunit Rpt3, a conserved residue among vertebrate, is phosphorylated during the G2/M phase to facilitate efficient protein translocation for degradation and cell proliferation, while phosphorylation of S361 of Rpn1 and S16 of α5 impair 26S proteasome assembly and cell proliferation^7,8,61^. Recently, in response to DNA damage, phosphorylation of Rpn10-S266 alters the conformation of its two ubiquitin-interacting motif, resulting in reduced binding to ubiquitinated DNA repair proteins and subsequent degradation for efficient DNA repair^9^. Our study demonstrates that, phosphorylation of α subunit, PsmA-S200 in *Sa. islandicus* impairs 26S proteasome assembly, akin to the effect of Rpn1-S361 phosphorylation in eukaryotes^7^. We speculate that PsmA-T213 phosphorylation may disrupt its interaction with Pba1/2 homologs and impair its incorporation into the *de novo* α ring, an early step of 20S proteasome assembly. Similarly, the eukaryotic α5-S16 residue is buried at the 19S-20S interface and its phosphorylation only occur in the free form of α5 or in protein complexes too small to be proteasomes^8^, indicating that this phosphorylation also impair the early assembly of the 20S proteasome. Moreover, our current and previous phosphoproteomic analysis^43^ have identified phosphorylation of S46, S59 and S64 in PsmA, in addition to S200 and T213. The proximity of S59 and S65 to the conserved K68A suggests that they may directly impact the activation of the 20S proteasome by PAN. Therefore, it is likely that phosphorylation of different residues in PsmA confer various strategies for proteasome regulation, similar to those in eukaryotes.

Although phosphorylation occurs at multiple residues of archaeal PsmA, none of them are conserved in Sulfolobales as revealed by sequence alignment. In contrast, phosphorylated residues in several subunits of eukaryotic proteasomes are highly conserved^7,8,61^, suggesting that phosphorylation-mediated regulation of proteasomes in eukaryotes may already exist in FECA, the first eukaryotic common ancestor. However, phosphorylation on archaeal PsmA might vary among different species while still serving similar functions, implying an early form of proteasome regulation by post-translational modification.

We observed that Δ*ePK2* did not display any apparent differences with the wild type in terms of cell growth, morphology, and flow cytometry profile when not synchronized (Fig. 2A-D). However, synchronized Δ*ePK2* cells exhibited a deficiency in cell cycle progression deficiency with a lack of G1 peak, presumably resulted from elevated degradation of CdvBs at critical time of the cell cycle (M-D phases)(Fig. 2E, 5E and 5F). It is possible that in asynchronized cells, PsmA would be phosphorylated by other protein kinase(s) that are constantly expressed throughout the entire cell cycle, albeit less efficiently (Fig. S1). In agreement with this, our *in vitro* kinase assay showed that ePK1, another highly active kinase, was also able to phosphorylate PsmA at a similar extent to ePK2 (Fig. 5C). Furthermore, ePK2 contains multiple tetratricopeptide repeats (TPR) that could potentially recruit other protein kinases (or partners) through protein-protein interaction for phosphorylation of S46, S59 and S65 *in vivo* which are not directly targeted by ePK2^62^. We speculate that treatment of Sulfolobales cells with acetic acid would induce a quiescent stage during which protein activities are reduced. Under such conditions, the complementary phosphorylation by other kinases may not function efficiently, leading to the assembly of active proteasome and a failure in reducing cellular proteasome activity in M-D phases. It would be interesting to investigate which kinase phosphorylates these residues and how the phosphorylation works. However, the lack of G1 peak in synchronized Δ*ePK2* cells and uncontrolled DNA replication in the enlarged cells overexpressing ePK2 may also indicate a decoupling of cell division and the initiation of DNA replication. Therefore, exploring other targets of ePK2 and the proteasome or utilizing alternative synchronization methods, such as the baby machine method^63^ or treating with the translation inhibitor tetracycline^64,65^, could help elucidate the phenotypic discrepancy between synchronized and asynchronized cells.

To summarize, our study demonstrates that protein phosphorylation plays a vital role in regulating the cell cycle progression in archaea. Based on our findings, we propose a model depicting ePK2 phosphorylation on the proteasome α subunit for cell division regulation in Sulfolobales (Fig 9). Following G2 phase, the expression of ePK2 gradually increases in conjunction with cell division proteins. This leads to the inhibition of the proteasome activity through phosphorylation of the α subunit at S200 and T213 by ePK2. This dual phosphorylation events prevent the assembly of the 26S proteasome (S200 phosphorylation) and results in the formation of inactive 26S proteasome (T213 phosphorylation), ultimately leading to the stabilization of cell division proteins CdvB/CdvB1/CdvB2 at mid-cell. During D phase, the down-regulation of ePK2 allows for the assembly and activation of proteasomes for degradation of CdvB proteins. Furthermore, reduced transcription of CdvB genes by aCcr1 will also facilitate preventing the accumulation of CdvBs. The coordinated action results in the progression of cell division through ring constriction and membrane abscission. When ePK2 is overproduced during M phase, continuous inhibition of the proteasome leads to the accumulation of CdvB proteins, causing cells to enlarge and exhibit increased DNA contents due to decoupling between cell division and DNA replication. Future work will focus on how phosphorylation affects proteasome assembly and how the eukaryotic proteasome phosphorylation mechanisms evolved from archaea especially Asgard archaea.

**Figure 9.**
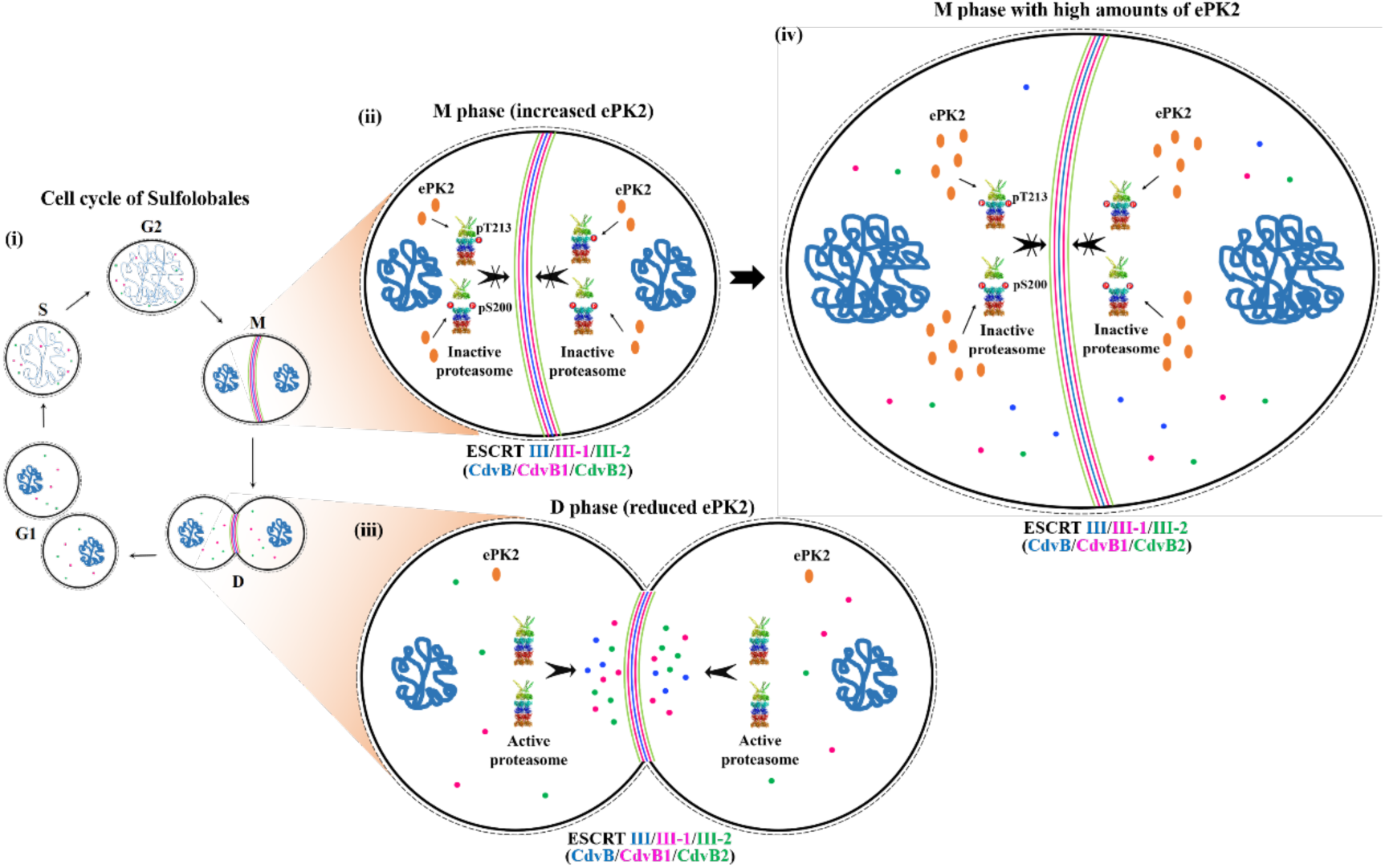
A proposed model of cell division regulation by ePK2 phosphorylation of the proteasome in Sulfolobales. (**i**) The cell cycle of Sulfolobales consists of G1, S, G2, M and D phases. (**ii**) Following the G2 phase, an increase in ePK2 expression leads to elevated PsmA phosphorylation. This results in the failure of 26S proteasome assembly (S200 phosphorylation) or the formation of inactive 26S proteasome (T213 phosphorylation). Inhibition of proteasome assembly halts the degradation of cell division proteins CdvB/CdvB1/CdvB2, allowing them to accumulate at the mid-cell. **(iii**) During the D phase, the expression of ePK2 decreases, facilitating the assembly and activation of proteasomes for the degradation of CdvB proteins. The reduced transcription of *cdvB* genes by aCcr1 (Yang *et al*., 2023) further stimulates the disassembly of CdvBs rings. This allows the cell cycle to progress with ring constriction and ultimately membrane abscission. (**iv**) When ePK2 is overproduced during M phase, the proteasome remains constantly inhibited, leading to the accumulation of CdvB proteins. This results in enlarged cells with increased DNA contents.

## Materials and methods

### Strains and growth conditions

The *pyrEF* and *lacS* deficient mutant *Sa. islandicus* REY15A (E233S) (hereafter E233S) and its derivatives were grown at 75 °C in STVU medium (basal salts supplemented with 0.2 % (w/v) sucrose (S), 0.2 % (w/v) tryptone (T), a mixed vitamin solution (V), and 0.01 % (w/v) uracil). The pH was adjusted to 3.3 with sulfuric acid as described previously^44^. E233S carrying the shuttle vector pSeSD with or without genes were grown in STV medium. The ATV was used for protein overexpression in *Sa. islandicus,* where sucrose was replaced with 0.2 % (w/v) D-arabinose (A). The solid medium was prepared using gelrite (0.8 % [w/v], Sigma, USA). Normally, cells were cultured to OD_600_=0.5–0.8, then transferred to fresh medium with an initial estimated OD_600_ of 0.05 for growth curve determination. PMSF was purchased from Sangon Biotech Co., Ltd. (Shanghai, China), protease inhibitor mixture and Suc-LLVY-AMC were purchased from MeilunBio (Dalian, China), and Bortezomib was purchased from Solarbio (Beijing, China).

### Construction of deletion, complementation and overexpression strains

Construction of the knockout and site-directed mutant strains was performed according to the endogenous CRISPR-Cas-based genome editing method as described previously^66^. The 40 nt DNA sequence downstream of CCA or TCA motif was selected as the protospacer and annealing of the two complementary oligonucleotides yielded a DNA fragment containing the designed spacer. The spacer fragment was inserted into the digested vector yielding the artificial mini-CRISPR plasmid. The donor DNA fragment was generated by splicing and overlapping extension (SOE)-PCR with *ApexHF* HS DNA Polymerase FS (Accurate Biotechnology, Hunan, China). Then the donor DNA fragment was digested with *Sph*I and *Xho*I and inserted into mini-CRISPR plasmid to generate the genome editing plasmids. The cells of *Sa. islandicus* E233S were transformed with the plasmid by electroporation^67^. The in-frame deletion and site-directed mutant strains were screened and verified by PCR amplification using the primers check-F and check-R primers and subsequent sequencing of the PCR products. The gene editing plasmid was removed by counter selection with uracil and 5-fluoroorotic acid (5-FOA). The strains, plasmids and the primers used in this study are listed in Supplementary Table S1, Table S2 and Table S3, respectively.

To construct ePK2 complementation strain, the ePK2 gene together with its native promoter sequence (NP-ePK2) was amplified using the genomic DNA of *Sa. islandicus* E233S by PCR. The DNA fragment was digested and ligated into the vectors pSeSD^47^ at *Sph*I/*Sal*I site, yielding the expression plasmid pSe-NP-ePK2. The plasmid was transformed into *Sa. islandicus* Δ*ePK2* cells for ePK2 complementation.

To construct gene overexpression strains, the gene fragments for ePK2, PsmA, and the proteasome assembly factors SiRe_1163 and SiRe_1724 were amplified by PCR and the fragment containing mutant derivative was obtained by SOE-PCR. The resulting DNA fragments were digested and ligated into the vectors pET22b and pSeSD at *Nde*I/*Sal*I site, yielding the expression plasmids for WT and mutant proteins. The sequences of the plasmids were verified by DNA sequencing. The pET22b plasmids were transformed into *E. coli* BL21(DE3) cells for the expression of recombinant proteins. The pSeSD plasmids were introduced into the cells of *Sa. islandicus* E233S by electroporation, yielding *Sa. islandicus* strains for protein overexpression.

### Cell synchronization

Cell synchronization was performed as described previously^29^. Briefly, *Sa. islandicus* cells were grown to log-phase and transferred for 3–4 times for activation of the cells. Cell synchronization was conducted by adding 6 mM acetic acid into a culture with OD_600_∼0.2 for 6 h. Then the cells were washed with 20 mM sucrose for 3 times and re-incubated in preheated medium which was set as the start point (0 h). The cells were then released from G2 phase during which samples were taken for flow cytometry and Western blotting at 1 h interval or as specified time points.

### Microscopy and flow cytometry

Cell morphology was analyzed by microscopy under a NIKON TI-E inverted fluorescence microscope (Nikon, Kobe, Japan). Samples from three independent cultures were examined. For flow cytometry, the culture (0.3 mL) of *Sa. islandicus* were harvested and fixed with 70 % ice-cold ethanol (final concentration) and stored at 4 °C. After 12 h, the fixed cells were collected by centrifugation at 800 *g* for 20 min and washed twice with 1 mL of PBS buffer. The pellets were resuspended in 150 μL PBS buffer and stained with SUPER Green Ⅰ (Fanbo Biochemicals, Beijing, China). The stained samples were analyzed by flow cytometry using the ImageStreamX MarkII Quantitative imaging system (Merck, Darmstadt, Germany) equipped with a 488 nm laser. A dataset of at least 20,000 cells was obtained for each sample. For synchronized cell samples, propidium iodide (Sigma-Aldrich) was used for DNA staining.

### Quantitative reverse transcription PCR (RT-qPCR)

Cells were collected by centrifugation at 6,000 *g* for 5 min and 1 mL of the SparkZol Reagent (SparkJade, Shandong, China) was added. Total RNA was extracted according to the manufacturer’s instructions. The first-strand cDNA was synthesized by reverse transcription of RNA using Evo M-MLV RT Mix Kit (Accurate Biotechnology, Hunan, China) according to the manufacturer’s protocol. The mRNA levels were evaluated by RT-qRCR using the SYBR Green Premix *Pro Taq* HS qPCR Kit (Accurate Biotechnology, Hunan, China) on the CFX96 Touch™ Real-Time PCR System (Bio-Rad, Hercules, CA, USA). The two-step PCR amplification standard procedure was as follows: 95 ℃ for 30 s, followed by 40 cycles of PCR that consisted of 95 ℃ for 5 s and 60 ℃ for 30 s. The relative transcription level of the target gene was calculated using the comparative threshold cycle (CT) method (2^−ΔΔCT^)^68^, and the TBP or 16S rRNA gene was used as reference. The qPCR primers used in this study are listed in Supplementary Tables S3.

### Western blot analysis

An aliquot of cell pellets was resuspended in a 100 μL of Buffer A (50 mM Tris-HCl pH 8.0, 200 mM NaCl, and 5% glycerol) and mixed with 5×SDS-PAGE loading buffer for SDS-PAGE analysis. The proteins in the PAGE gel were transferred onto a PVDF membrane at 200 mA for 2 h at 4 ℃. The membrane was washed and incubated with a primary antibody (Mabnus, Wuhan, China) and then the secondary anti-rabbit HRP-conjugate antibody (Kermey, Zhengzhou, China) following the standard protocol for Western blotting. The band was visualized with Immobilon^TM^ Western Chemiluminescent HRP Substrate (Kermey, Zhengzhou, China) and the image was obtained by Imagequant^TM^ 400 (GE Healthcare, UK). The ePK2, PsmA and PsmB1 specific antibodies were prepared in rabbits using full length proteins expressed in and purified from *E. coli*. Phosphorylated protein was detected with Phospho-Threonine/Tyrosine antibody (#9381S, Cell signaling technology, MA, USA).

### Protein purification

The pET22b plasmids carrying ePK2 or PsmA genes were transformed into *E. coli* BL21 (DE3)-CodonPlus-RIL for protein expression. The procedure for protein induction and purification in *E. coli* cells was the same as previously described^45^. Briefly, after induction, the cells were harvested and resuspended in buffer A for lysis by sonication. The soluble fractions were heated at 70 ℃ for 30 min and after centrifugation of 10000 *g* for 20 min the supernatants were purified by a Ni-NTA column and Superdex^TM^ 200 10/300 column subsequently (GE Health, UK). The protein concentration was determined by the Bradford method with bovine serum albumin (BSA) as the standard.

For the purification of proteasomes from various PsmA overexpression strains, 3 L of each sample (OD_600_∼0.4) cultivated in STV medium were collected and resuspended in buffer A for lysis by sonication. The soluble fractions were subjected for purification using a Ni-NTA column and Superdex^TM^ 200 10/300 column. The fractions obtained by gel filtration were analyzed by SDS-PAGE, Western blotting and focused ion beam electron microscopy.

### Phosphoproteomic analysis

Sis/pSeSD and Sis/pSeSD-ePK2 were cultured in ATV medium to OD_600_ = 0.5–0.6. The cells were collected by centrifugation at 6,000 *g* for 10 min and washed with PBS buffer (137 mM NaCl, 2.7 mM KCl, 10 mM Na_2_HPO_4_, 2 mM KH_2_PO_4_) for phosphoproteomic analysis by Bioprofile (Shanghai, China).

The cell pellets were resuspended in lysis buffer (4% SDS, 100 mM DTT, 150 mM Tris-HCl pH 8.0). The samples were boiled for 3 min and further sonicated for 2 min. Undissolved cellular debris were removed by centrifugation at 16000 rpm for 15 min. The supernatant was collected and quantified with a BCA Protein Assay Kit (Bio-Rad, USA). Digestion of protein was performed according to the FASP procedure described by Wisniewski et al.^69^. Briefly, 1 mg total protein was mixed with 10 mM DTT and boiled for 5 min. After addition of 50 mM iodoacetamide (in 8 M Urea, 150 mM Tris-HCl pH 8.0), the mixture was incubated for 1 min at 600 rpm followed by 30 min at RT in darkness. Six volumes of cold acetone was added and incubated at -20 ℃ overnight for protein precipitation. After centrifugation at 16000 *g* for 15 min, the protein pellet was washed with acetone for twice and dried in a fune hood. The protein suspension was digested in 150 μL trypsin buffer (Promega) (200 μg trypsin in NH_4_HCO_3_ buffer) overnight at 37 ℃. Then the peptides were acidified with 0.1% trifluoroacetic acid (TFA), desalted with C18 cartridge (Thermo, USA) and vacuum dried. The dried peptides were dissolved in 0.1% TFA and desalted by Thermo desalting spin column. Finally, the concentrations of peptides were determined with OD_280_ by Nanodrop device (Thermo, USA).

Peptides were labeled with TMT reagents according to the manufacturer’s instructions (Thermo Fisher Scientific). Each aliquot (200 μg of peptide equivalent) was reacted with one tube of TMT reagent, respectively. The labeled peptides were mixed equally, desalted by Peptide desalting spin column (Thermo Fisher Scientific) and vacuum dried. For phosphopeptide enrichment, Fe-NTA Phosphopeptide Enrichment kit (Thermo. A32992) were used for digested peptide mixtures from each sample according to the manufacturer’s instructions. The eluted phosphopeptides were vacuum-dried immediately for fractionation by high pH reverse-phase with STAGETip C18. The total of 6 phosphopeptides fractions were suspended with 0.1% formic acid (FA) for further LC-MS/MS analysis.

LC-MS analysis was performed on an Easy nLC 1200 chromatographic system (Thermo Fisher Scientific). For phosphoproteomic study, the phosphopeptides were loaded to Trap column (100 μm × 20 mm; 5 μm ReproSil-Pur C18 beads, Dr. Maisch GmbH, Ammerbuch, Germany) followed by a self-packed column (75 μm × 150 mm; 3 μm ReproSil-Pur C18 beads, Dr. Maisch GmbH) using buffer A (0.1% formic acid) for gradient separation at a flow rate of 300 nL/min. Peptide were eluted as following procedure: 0–2 min, 2%–8% linear Buffer B (0.1% formic acid and 85% acetonitrile); 2–45 min, 8%–28% linear Buffer B; 45–50 min, 28%–40% linear Buffer B; 50–52 min, 40%–100% linear Buffer B; 52–60 min, 100% Buffer B. Fractioned peptides were applied to a Q-Extractive HF-X mass spectrometer for 60 min DDA analysis. For MS data acquisition, full MS scans were surveyed from m/z 350 to m/z 1800 at a resolution of 60,000 at m/z 200 with an AGC target values of 3e6 and a maximum injection time 50 ms. Then data-Dependent top 20 MS/MS scans were applied by higher-energy collision dissociation (HCD) with normalized collision energy 32 at a resolution of 45,000 at m/z 200 with an AGC target values of 1e5 and a maximum injection time 50 ms. The isolation window was set to 1.2 m/z.

The phosphoproteome data were imported into Proteome Discoverer 2.4 for data interpretation and protein identification against the *Sa. islandicus* (strain REY15A) database from Uniprot (downloaded on 01/10/2022, and including 2631 protein sequences), which is sourced from the protein database at https://www.uniprot.org/uniprot/. For phosphoproteome MS spectra were searched with 4.5 ppm mass tolerance for precursor ions and 20 ppm mass tolerance for fragment ions, fully Tryptic restriction and maximal two missed cleavage sites. The modifications were variable phosphorylation (+79.96633) on serine (S), threonine (T) and tyrosine (Y). The search results were filtered and exported with <1% false discovery rate (FDR) at site level, peptide-spectrum-matched level, and protein level, respectively. MaxQuant analysis was filtered only for those phosphorylation sites that were confidently localized (class I, localization probability > 0.75).

For phosphoproteomic data, the t-test (P value) analysis and fold change (FC) was followed and used to determine statistical significance (Q < 0.2) for each comparison. Expression data were grouped together by hierarchical clustering according to the peptide level. To annotate the sequences, information was extracted from Kyoto Encyclopedia of Genes and Genomes (KEGG), and Gene Ontology (GO). GO and KEGG enrichment analyses were carried out with the Fisher’s exact test, and FDR correction for multiple testing was also performed.

### *In vitro* kinase activity assay

For the *in vitro* phosphorylation activity assay of the protein kinases on PsmA (or its mutants), indicated amounts of protein kinases and PsmA were added into a reaction mixture (20 μL) containing 25 mM Tris-HCl pH 8.0, 5 mM NaCl, 5 mM MgCl_2_, 1 mM DTT, and 1 mM ATP). The mixture was incubated at 65 ℃ for 30 min and the reaction was stopped by adding 5×SDS-PAGE loading buffer and boiling for 10 min. The samples were analyzed by 15% SDS-PAGE and Western blotting with Phospho-Threonine/Tyrosine antibody. For the kinase assay using radiolabeled [γ-^32^P] ATP, 0.025 μCi [γ-^32^P] ATP (111 TBq/mmol, Hartmann Analytic, Germany) was used in the reactions. The autoradiographs were quantified by the software ImageQuant 5.2.

### Focused ion beam electron microscopy

The protein particles in the PsmA samples purified from *Sa. islandicus* were analyzed by focused ion beam electron microscopy. The protein samples (1–1.5 mg/mL) were dropped onto a carbon film supported copper mesh and stained with 1% acetic acid for 30 s. After dried by an infrared lamp, the samples were observed under a ZEISS crossbeam 550 electron microscope in the STEM model.

### Sucrose density gradient fractionation

For detection of proteasome oligomerization and association profiles, a volume of 20 mL *Sa. islandicus* cells (OD_600_∼0.3–0.4) was collected and resuspended in 1 mL of Buffer A. After sonication, the samples were centrifuged at 20,000 *g* for 30 min at 4 ℃. The supernatants (0.5 mL) were layered on top of 12 mL of 5–25% (or 5–20% as indicated) sucrose gradients made in a buffer (containing 50 mM Tris-HCl pH 7.5, 150 mM NaCl, 10% glycerol). Separation of the gradients was performed in a centrifuge with a Beckman SW41 rotor for 20 h at 35000 rpm at 4 ℃. The gradients were fractionated from the top of the sucrose mixture into 500 μL aliquots. Aliquots (20 μL) of the fractions were analyzed by Western blotting with PsmB1 antibody. All experiments were repeated at least three times.

### Proteasome activity assay

The proteasome activity of whole cell lysis was evaluated by the Fluorescence substrate Suc-Leu-Leu-Val-Try-AMC (Macklin, Shanghai, China). A volume of 20 mL *Sa. islandicus* cells (OD_600_∼0.3–0.4) was collected and resuspended in 1 mL of Buffer A for lysis by sonication. The concentrations of total proteins in the soluble fractions were determined by the Bradford method. An aliquot of 0.5 μg total protein samples was applied for the proteasome activity assay in a 100 μL reaction containing 50 mM Tris-HCl pH 7.5, 250 mM sucrose, 5 mM MgCl_2_, 1 mM DTT, 2 mM ATP, and 20 μM Suc-LLVY-AMC. The reaction mixture was incubated at 70 ℃ for 30 min. The fluorescence signals were detected by PerkinElmer enspire microplate spectrometer with excitation at 345 nm and emission at 445 nm (CT, USA).

### Protein structure prediction

The monomers and α ring structures of wild type PsmA, S200D, and T213E were predicted by Alphafold server (https://alphafoldserver.com/about). The structures were aligned and analyzed using the software PyMOL.

## Data availability

All data supporting the findings of this study are available within the article and its Supplementary Information, or from the corresponding author upon reasonable request. The mass spectrometry proteomics data have been deposited to the ProteomeXchange Consortium via the PRIDE partner repository with the dataset identifier PXD061899.

## Acknowledgements

We would like to thank the former lab members, Fan Zhou and Dr. Junfeng Liu, for initial work on this project. We are grateful to Prof. Buzz Baum at MRC laboratory for insightful suggestions. Thanks all the lab members of the CRISPR and Archaea Biology Research Centre for helpful discussions and technicians from the Core Facilities for Life and Environmental Sciences, State Key Laboratory of Microbial Technology of Shandong University for assistance.

