## Supplementary Figures and Tables for "Phosphorylation of the α subunit inhibits proteasome assembly and regulates cell division in an archaeon"

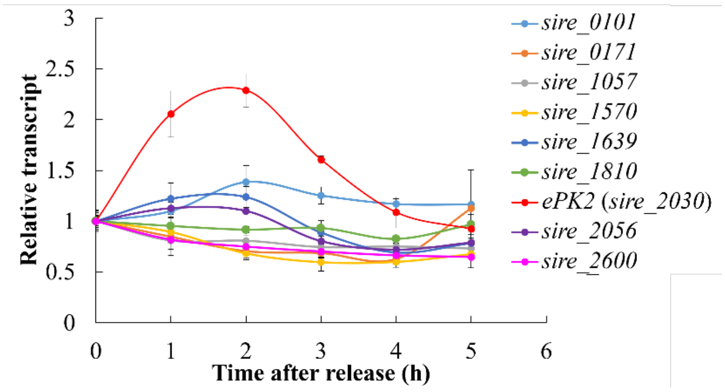


**Figure S1. Transcriptional profiles of synchronized *Sa. islandicus* cells.** Data were extracted from Yang *et al.,* published in Nucleic Acids Research in 2023. The cells were synchronized by acetic acid treatment for 6 h and released by washing with 20 mM sucrose and culturing in fresh preheated medium. Samples were collected at the indicated time points (at 1 to 5 h) after release. Relative transcripts were calculated as FPKM (Fragments Per Kilobase transcript per Million mapped reads) of each gene divided by the level at 0 h, which was set as 1.

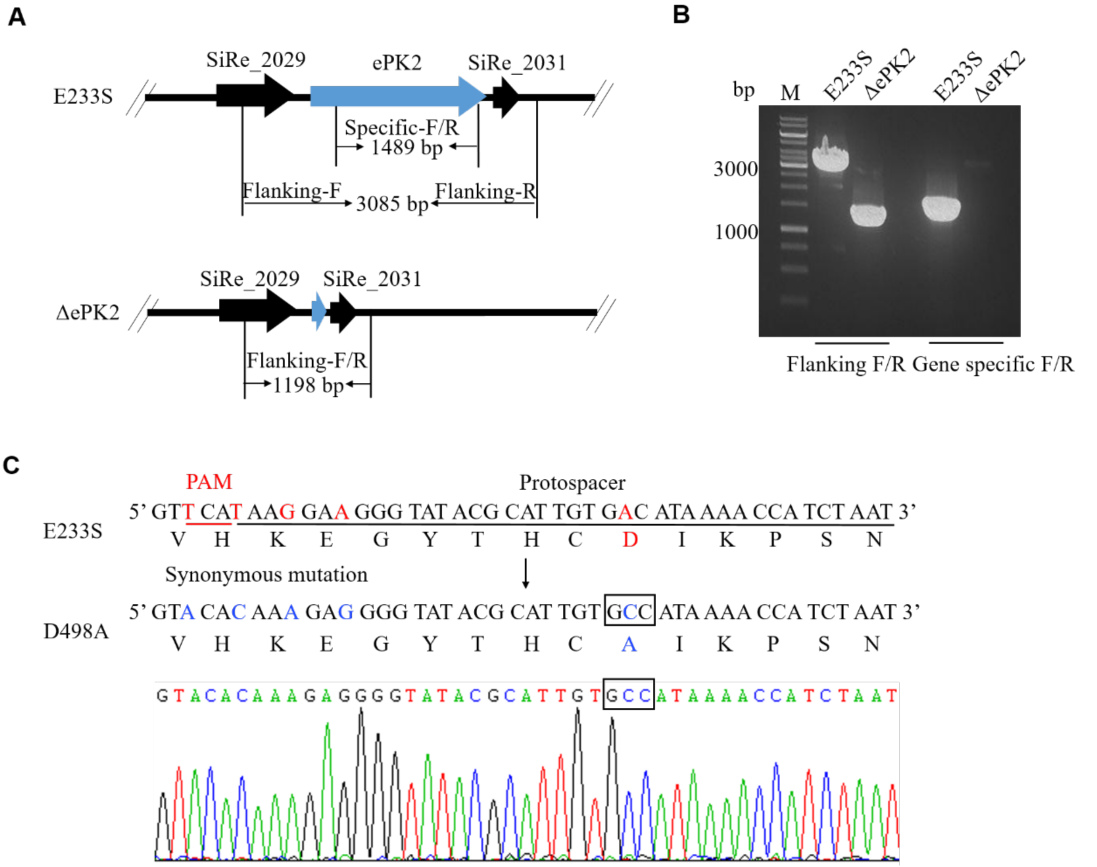


**Figure S2. Validation of genome editing in Δ*ePK2* and ePK2-D498A.** (**A**) Schematic depiction of PCR verification for the ePK2 deletion strain (Δ*ePK2*) with E233S serving as the wild-type control. (**B**) Verification of *ePK2* deletion by PCR using the flanking and specific primers. (**C**) Verification of *in situ* mutant strains ePK2-D498A through sequencing. Upper panels show the mutagenic strategies with red indicating bases to be mutated and blue indicating the mutated bases at the corresponding loci of the mutant strains. Synonymous mutations on the PAM and the protospacer sequence were introduced to inactivate Type I-A DNA interference. The target mutations are indicated in rectangles. Lower panels display chromatographs of the sequencing results for the single transformants.

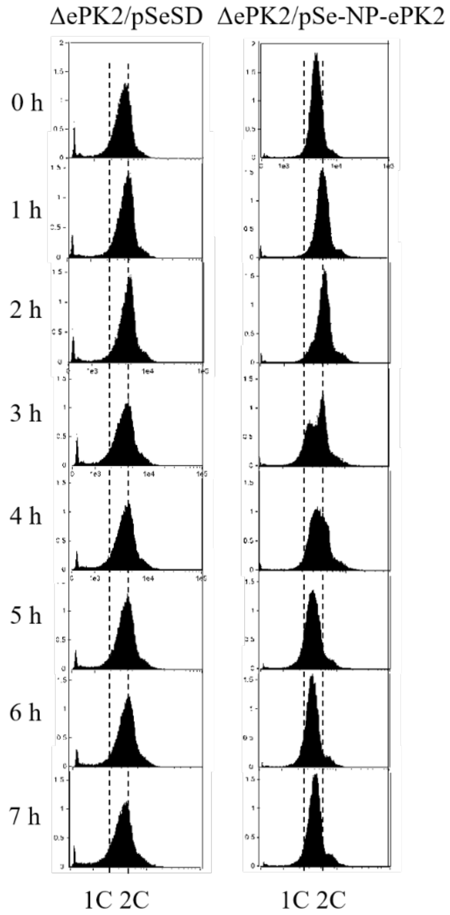


**Figure S3. G1 peak recovered after complementation of ePK2 in synchronized Δ*ePK2*.** Δ*ePK2* cells carrying pSeSD or pSe-NP-ePK2 were synchronized by acetic acid treatment at $OD_{600}$~0.2. Cells were released by washing out acetic acid, and cultivated in preheated fresh medium. Samples were taken at the indicated time points for flow cytometry analysis.

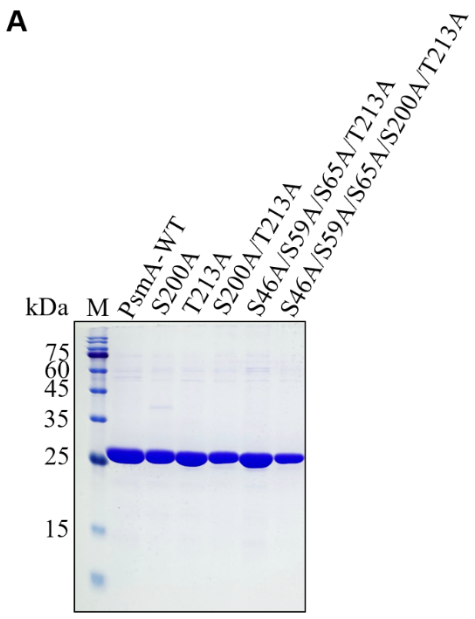


**Figure S4. SDS-PAGE analysis of the purified proteins.** (**A**) PsmA and its mutants. (**B**) Protein kinases and its mutants. The protein were purified as described in the Method and Materials. Briefly, total cell extracts were heat-treated by 70ºC for 30 min. The soluble proteins were purified using a Ni-NTA column and subsequent Superdex$^{TM}$ 200 Increase column (GE). WT, wild type; TM, transmembrane.

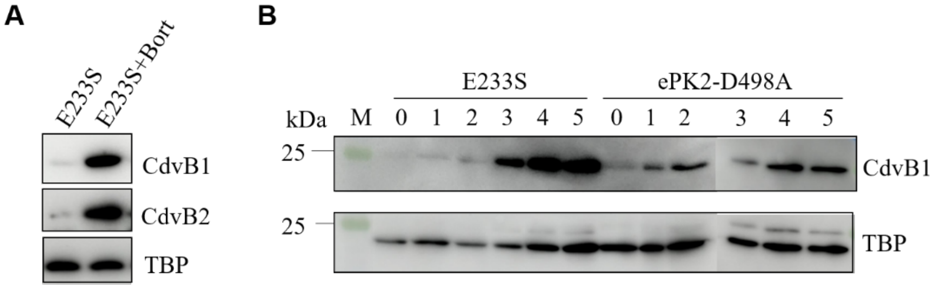


**Figure S5. Detection of cell division proteins in Bortezomib-treated E233S cells and synchronized ePK2-mutant cells.** (**A**). Detection of CdvB1 and CdvB2 in cells treated with Bortezomib. The cells were cultured to the early log phase and treated or mock-treated with the proteasome specific inhibitor Bortezomib (+Bort) for 6 h before sample collection. (**B**) Detection of CdvB1 in cells of synchronized ePK2-mutant strain. The wild type E233S and the ePK2-mutant strains, ePK2-D498A, were synchronized by acetic acid treatment at $OD_{600}$~0.2. Cells were released by washing out acetic acid and inoculating into fresh preheated medium. Samples were taken at the indicated time points. Western blotting analysis was performed with protein specific antibodies as indicated with TBP used as the loading control.

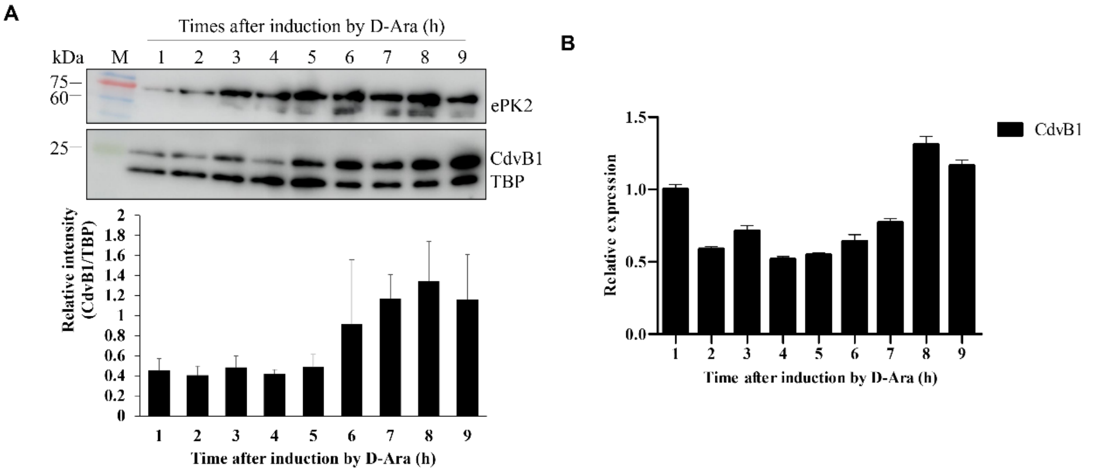


**Figure S6. Detection of the protein levels of CdvB1 and ePK2 (A) and the transcription level of *cdvB1* in ePK2 overexpression cells (B).** The ePK2 overexpression strain was cultured to early log phase ($OD_{600}$~0.2) and ePK2 was induced by the addition of D-arabinose (D-Ara). Samples were taken at specified time points after induction. Western blotting analysis was performed with specific antibodies as indicated. Total mRNA of the samples was extracted for RT-qPCR with *tbp* gene as a reference.

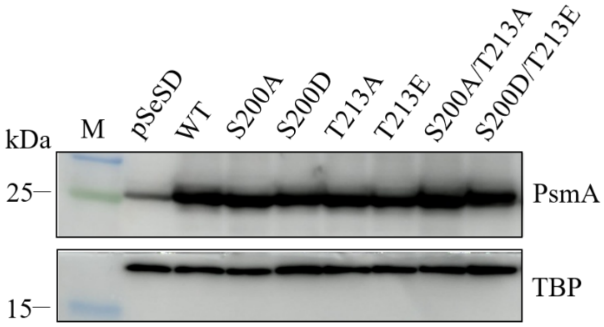


**Figure S7. Detection of PsmA protein levels in the overexpression strains of PsmA and its mutants by Western blotting.** Cultures were grown to the early log phase ($OD_{600}$~0.2–0.3) before samples were taken. Western blotting was performed using anti-PsmA antibody with TBP used as a control.

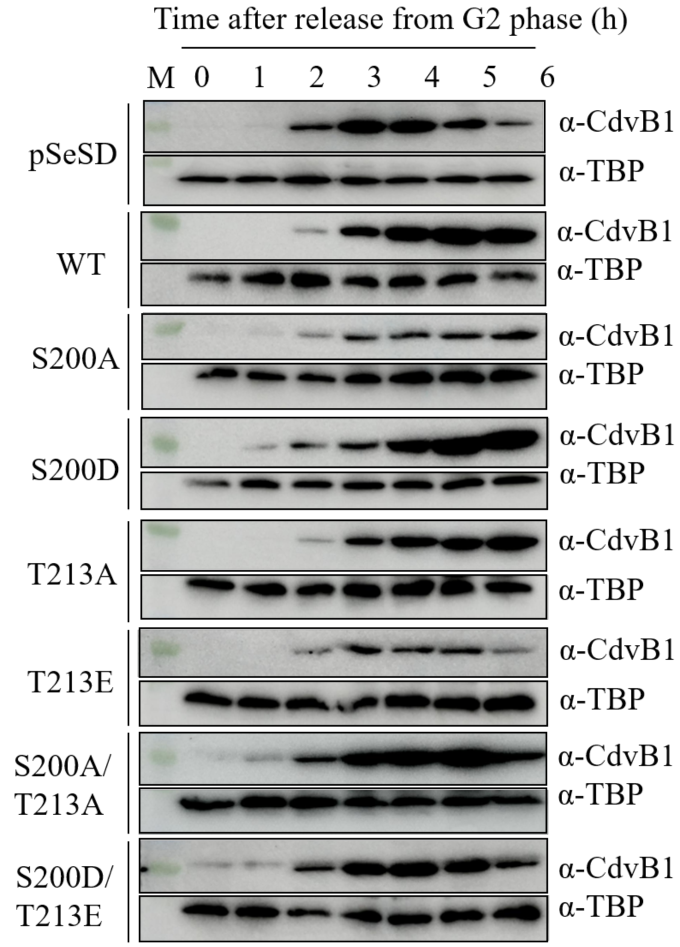


**Figure S8. Overexpression of mutants T213E and S200D/T213E did not inhibit CdvB1 degradation.** Overexpression strains of PsmA and its mutants were cultured in STV medium and synchronized by acetic acid treatment at $OD_{600}$~0.2 at which arabinose was added for protein induction. The cell cycle was released by washing out acetic acid and inoculating into fresh preheated ATV medium. Samples were taken at specified time points and analyzed by Western blotting with CdvB1 and TBP antibodies.

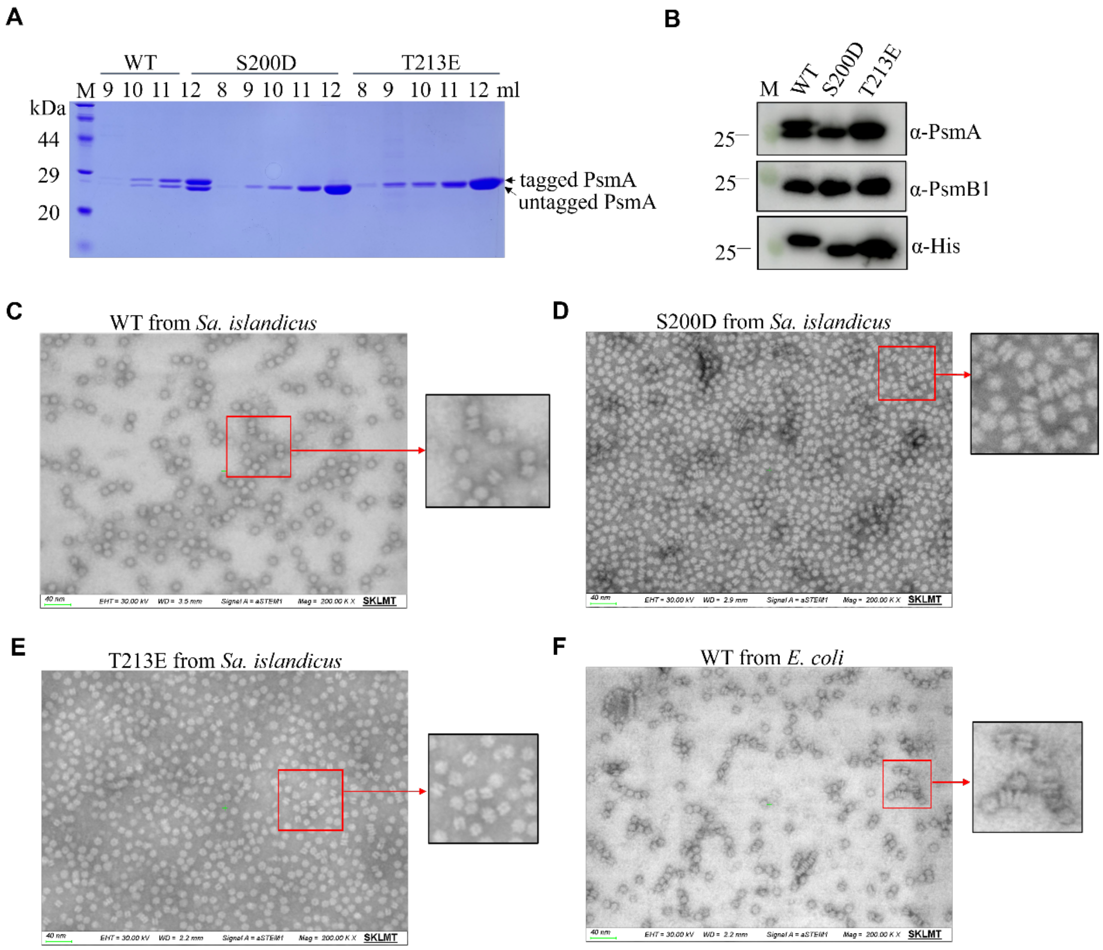


**Figure S9. Analysis of proteasomes by electron microscopy.** (**A**) Purification of proteasomes from PsmA, S200D, and T213E overexpression strains. The strains were cultured in STV medium (3 L) and cells were collected and lysed by sonication. The soluble fractions were subjected to protein purification by Ni-NTA and gel filtration. The fractions containing PsmA proteins were analyzed by SDS-PAGE. The numbers indicate the fractions in gel filtration. (**B**) Detection of PsmA and PsmB1 in the purified PsmA samples by Western blotting. The gel filtration fraction samples were analyzed by Western blotting with PsmA and PsmB1 specific antibodies. (**C-E**) Observation of the purified proteasomes by electron microscopy. The purified PsmA (C), S200D (D) and T213E (E) from cells of *S. islandicus* strains and PsmA from *E. coli* (F), were observed by Focused Ion Beam Scanning Electron Microscope. The images in red rectangles were zoomed in at the right panels. Scale bar (green), 40 nm.

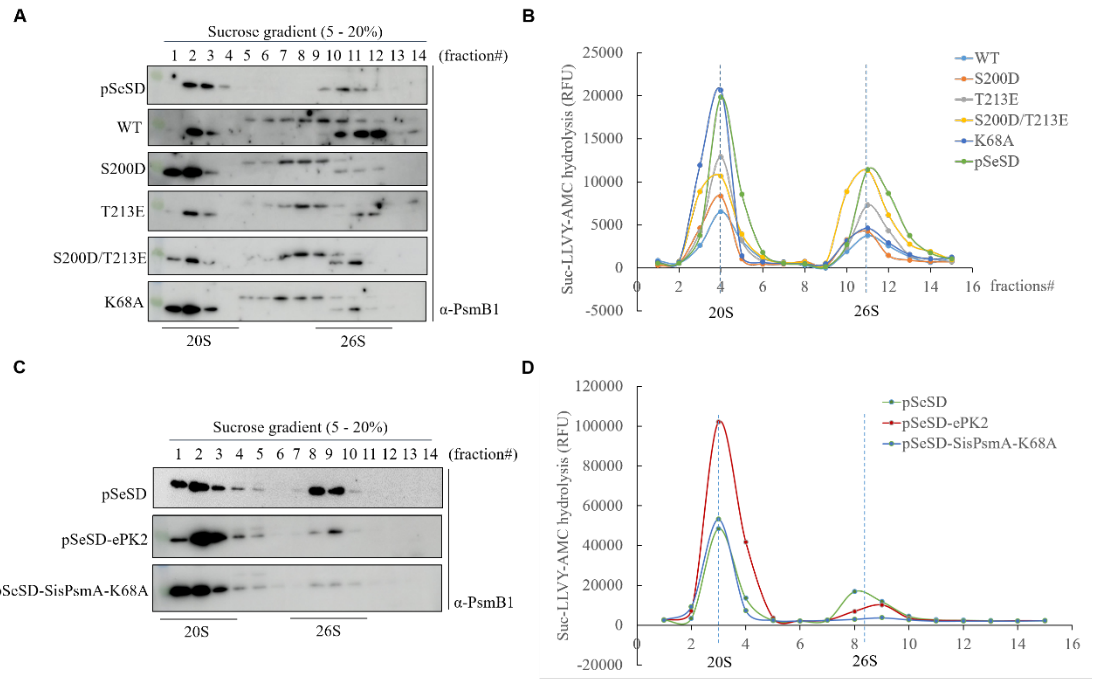

**Figure S10. Overexpression of PsmA(K68A) disrupts 26S assembly, while overexpression of S200D/T213E does not affect cellular 26S assembly. (A)** Analysis of 20S and 26S proteasomes in PsmA overexpression strains. A 0.5 ml aliquot of soluble whole cell lysate (WCL) from PsmA overexpression strains was centrifuged in sucrose gradients ranging from 5%-25% Samples were collected at 1 ml intervals and analyzed by Western blotting with anti-PsmB1 antibody. The positions of the 20S and 26S proteasomes are indicated at the bottom of the figure. (**B**) Proteasome activities of pSeSD-PsmA (WT), S200D, T213E, S200D/T213E, K68A, and the control pSeSD WCLs. The positions of 20S and 26S proteasomes are indicated at the bottom of the curves. (**C)** and (**D**) Analysis of the 20S and 26S proteasomes in ePK2 overexpression strain by sucrose density gradient fractionation and the proteasome activity assay.

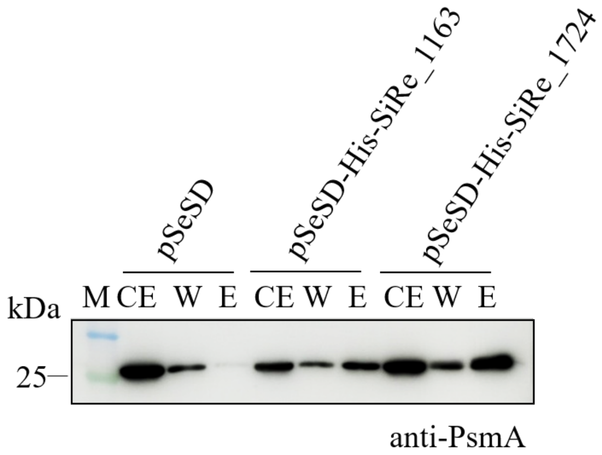


**Figure S11. Western blotting analysis to detect the interaction between PsmA and the putative proteasome assembly factors, SiRe_1163 and SiRe_1724 by *in vivo* pull-down.** The overexpression strains, E233S/pSeSD-His-SiRe_1163 and E233S/pSeSD-His-SiRe_1724, were cultivated in 300 mL STV medium. His-tagged proteins were purified with Ni-NTA columns. Samples of whole cell extracts (CE), wash fractions (W) (with 30 mM imidazole) and eluted fractions (E) (with 300 mM imidazole) were analyzed by Western blotting using anti-PsmA antibody. The strain with the empty vector pSeSD was used as a control.

### Supplementary Tables

**Table S1.** Strains used in this study

| Strain | Properties | Source or reference |
|---|---|---|
| *S. islandicus* E233S | *S. islandicus* REY15A Δ*pyrEF* Δ*lacS* | (Deng *et al.*, 2009) |
| Δ*ePK2* | *SisePK2* knockout in E233S | this study |
| Δ*ePK2*/pSe-NP-ePK2 | Complementation of ePK2-His in Δ*ePK2* | |
| ePK2-D498A | ePK2 with an in-frame mutation D498A in E233S | this study |
| E233S/pSeSD-ePK2-His | ePK2-His overexpression in E233S | this study |
| E233S/pSeSD-ePK2-D498A-His | ePK2-D498A-His overexpression in E233S | this study |
| E233S/pSeSD-ePK2-K393A/D498A-His | ePK2-K393A/D498A-His overexpression in E233S | this study |
| E233S/pSeSD-PsmA-His | PsmA-His overexpression in E233S | this study |
| E233S/pSeSD-PsmA-S200A-His | PsmA-S200A-His overexpression in E233S | this study |
| E233S/pSeSD-PsmA-S200D-His | PsmA-S200D-His overexpression in E233S | this study |
| E233S/pSeSD-PsmA-T213A-His | PsmA-T213A-His overexpression in E233S | this study |
| E233S/pSeSD-PsmA-T213E-His | PsmA-T213E-His overexpression in E233S | this study |
| E233S/pSeSD-PsmA-S200A/T213A-His | PsmA-S200A/T213A-His overexpression in E233S | this study |
| E233S/pSeSD-PsmA-S200D/T213E-His | PsmA-S200D/T213E-His overexpression in E233S | this study |
| E233S/pSeSD-PsmA-K68A-His | PsmA-K68A-His overexpression in E233S | this study |
| E233S/pSeSD-His-SiRe_1163 | His-SiRe_1163 overexpression in E233S | this study |
| E233S/pSeSD-His-SiRe_1724 | His-SiRe_1724 overexpression in E233S | this study |

**Table S2.** Vectors used in this study

| Vectors | Properties | Source or reference |
|---|---|---|
| pGE | *Sa. islandicus*-*E. coli* shuttle vector containing mini-CRISPR and *pyrEF* for CRISPR-Cas based gene editing | (Li *et al.*, 2016) |
| pGE-*ePK2*-KO | gRNA targeted *ePK2* knock out vector | this study |
| pSeSD | *Sa. islandicus* expression vector containing multiple cloning sites and two sequences for 6×His tag at both the 5' and 3' of the cloning sites | (Peng *et al.*, 2012) |
| pSeSD-ePK2-His | pSeSD containing C-terminal 6×His tagged *Sis*ePK2 | this study |
| pSe-NP-ePK2-His | pSeSD containing C-terminal 6×His tagged *Sis*ePK2 with its native promoter replacing the araS promoter | this study |
| pSeSD-ePK2-D498A-His | pSeSD containing C-terminal 6×His tagged ePK2-D498A | this study |
| pSeSD-ePK2-K393A/D498A-His | pSeSD containing C-terminal 6×His tagged ePK2-K393A/D498A | this study |
| pSeSD-PsmA-His | pSeSD containing C-terminal 6×His tagged PsmA | this study |
| pSeSD-PsmA-S200A-His | pSeSD containing C-terminal 6×His tagged PsmA-S200A | this study |
| pSeSD-PsmA-S200D-His | pSeSD containing C-terminal 6×His tagged PsmA-S200D | this study |
| pSeSD-PsmA-T213A-His | pSeSD containing C-terminal 6×His tagged PsmA-T213A | this study |
| pSeSD-PsmA-T213E-His | pSeSD containing C-terminal 6×His tagged PsmA-T213E | this study |
| pSeSD-PsmA-S200A/T213A-His | pSeSD containing C-terminal 6×His tagged PsmA-S200A/T213A | this study |
| pSeSD-PsmA-S200D/T213E-His | pSeSD containing C-terminal 6×His tagged PsmA-S200D/T213E | this study |
| pSeSD-PsmA-K68A-His | pSeSD containing C-terminal 6×His tagged PsmA-K68A | this study |
| pSeSD-His-SiRe_1163 | pSeSD containing N-terminal 6×His tagged SiRe_1163 | this study |
| pSeSD-His-SiRe_1724 | pSeSD containing N-terminal 6×His tagged SiRe_1724 | this study |
| pET22b | *E. coli* expression vector containing the T7 promoter and the sequence for 6×His tag at the 3' of the cloned gene | Novagen |
| pET22b-ePK1-His | pET22b containing C-terminal 6×His tagged ePK1 | (Huang *et al.*, 2017) |

| | | |
|---|---|---|
| pET22b-ePK2-His | pET22b containing C-terminal 6×His tagged ePK2 | this study |
| pET22b-ePK2-D498A-His | pET22b containing C-terminal 6×His tagged ePK2-D498A | this study |
| pET22b-PsmA-His | pET22b containing C-terminal 6×His tagged PsmA | this study |
| pET22b-PsmA-S200A-His | pET22b containing C-terminal 6×His tagged PsmA-S200A | this study |
| pET22b-PsmA-T213A-His | pET22b containing C-terminal 6×His tagged PsmA-T213A | this study |
| pET22b-PsmA-S200A/T213A-His | pET22b containing C-terminal 6×His tagged PsmA-S200A/T213A | this study |
| pET22b-PsmA-S46A/S59A/S65A/T213A-His | pET22b containing C-terminal 6×His tagged PsmA-S46A/S59A/S65A/T213A | this study |
| pET22b-PsmA-S46A/S59A/S65A/S200A/T213A-His | pET22b containing C-terminal 6×His tagged PsmA-S46A/S59A/S65A/S200A/T213A | this study |

**Table S3.** Primers used in this study

| Primers | Sequence (5’-3’)* |
|---|---|
| SisePK2KO-Spacer-F | AAGTAAGGAAGGGTATACGCATTGTGACATAAAACCATCTAAT |
| SisePK2KO-Spacer-R | AGCATTAGATGGTTTTATGTCACAATGCGTATACCCTTCCTTA |
| SisePK2KO-L-arm-*Sph*I-F | ATTC**GCATGC**TTTAGAAGAGGCCATGTC |
| SisePK2KO-L-R-arm SOE-R | CTAACTTCGCCCTTTAGCACATTAACTCTGTTC |
| SisePK2KO-L-R-arm SOE-F | GAACAGAGTTAATGTGCTAAAGGGCGAAGTTAG |
| SisePK2KO-R-arm-*Xho*I-R | TTGG**CTCGAG**GCTAGATGACGTTGTAATG |
| SisePK2-Flanking-F | GGTAGATCAAGGCATTGTAAATTGTATTGG |
| SisePK2-Flanking-R | CGTATCTGGACAAATCCCAGTAGATCCT |
| ePK2-D498A-Larm-SphI-F | ATAG**GCATGC**GAGCCCTTTGATGCTGCA |
| ePK2-D498A-LR-arm-SOE-R | CAATGCGTATACCCCTCTTTGTGTACTTCTATAACGGC |
| ePK2-D498A-LR-arm-SOE-F | AAAGAGGGGTATACGCATTGTGCCATAAAACCATCTAAT |
| ePK2-D498A-Rarm-XhoI-R | TTGC**CTCGAG**CTAACTTCGCCCTTTAG |
| ePK2-D532-spacer-F | AAG GATCTAGGTTCATCGGTAAAAATAGGTACTCCAGTAATGC |
| ePK2-D532-spacer-R | AGC GCATTACTGGAGTACCTATTTTTACCGATGAACCTAGATC |
| ePK2-D532A-LR-arm-SOE-R | GACCCAAGAGCACTTAGTTTAGGTACAACTTCAA |
| ePK2-D532A-LR-arm-SOE-F | AGTGCTCTTGGGTCATCGGT AAAAATAGGTAC |
| SisePK2-NdeI-F | CGGG**CATATG**GTGCAGTTAGTATTACAGTT |
| SisePK2-SalI-NoStop-R | GTGGCTG**GTCGAC**ATAGCTTATAAGCTT |
| NP-ePK2-SphI-F | ACAT**GCATGC**AATAACTTGGAAACTTTGAAATGAATAGAAACC |
| NP-ePK2-SalI- NoStop-R | TTGAC**GTCGAC**ATAGCTTATAAGCTTTTCTGCTTCACTG |
| SisePK2-D498A-R | GGGTATACGCATTGTGCCATAAAACCATCT |
| SisePK2-D498A-F | AGATGGTTTTATGGCACAATGCGTATACCC |
| SisePK2-D532A-R | CCGATGAACCTAGAGCAGATAGTTTAGGTA |
| SisePK2-D532A-F | TACCTAAACTATCTGCTCTAGGTTCATCGG |
| SisPsmA-NdeI-F | GGATTTG**CATATG**GCGTTCGGACCAGCC |
| SisPsmA-SalI-R | GGAG**GTCGAC**TAACTTCTGTAAATACATATTC |
| SisPsmA-S46A-R | GCTTGCAATAACAACACCAGCTTTTGATTTAATAC |
| SisPsmA-S46A-F | GTATTAAATCAAAAGCTGGTGTTGTTATTGCAAGC |

| | |
|---|---|
| SisPsmA-S59A-R | ACTATCTACATCTAATAATTCTTGGGCTTTTCTTTT |
| SisPsmA-S59A-F | AAAAGAAAAGCCCAAGAATTATTAGATGTAGATAGT |
| SisPsmA-S65A-R | TACTTTCTCTATAGCATCTACATCTAATA |
| SisPsmA-S65A-F | TATTAGATGTAGATGCTATAGAGAAAGTA |
| SisPsmA-K68A-R | CGATTAAAAATACTGCCTCTATACTATCT |
| SisPsmA-K68A-F | AGATAGTATAGAGGCAGTATTTTTAATCG |
| SisPsmA-S200A-R | GCTTAAGAGTAGCCGCTAAGGCCTTTAAG |
| SisPsmA-S200A-F | CTTAAAGGCCTTAGCGGCTACTCTTAAGC |
| SisPsmA-S200D-R | GCTTAAGAGTAGCGTCTAAGGCCTTTAAG |
| SisPsmA-S200D-F | CTTAAAGGCCTTAGACGCTACTCTTAAGC |
| SisPsmA-T213A-R | CCTATTTCCACAGCATTTGGAGTTAACT |
| SisPsmA-T213A-F | AGTTAACTCCAAATGCTGTGGAAATAGG |
| SisPsmA-T213E-R | CCTATTTCCACTTCATTTGGAGTTAACT |
| SisPsmA-T213E-F | AGTTAACTCCAAATGAAGTGGAAATAGG |
| SiRe_1163-NdeI-NHis-F | TTATTCTCATATGCACCACCACCATCATCACGATGAATTGAGTGAAGC |
| SiRe_1163-SalI-Stop-R | CCTGGTCGACTCACGCATAGGTGTATGG |
| SiRe_1724-NdeI-NHis-F | TTATTCTCATATGCACCACCACCATCATCACAGTAATGTTAAGATAATAATAAAG |
| SiRe_1724-SalI-Stop-R | CCTGGTCGACTTACATATAGACTCTATCAGC |
| pGE-F | CCGAATTTATCTATCGCTTTTCTCTCTC |
| pGE-R | CGGACATATTTGCCCTAACAGATAAG |
| TBP-q-F | GTGGCAACAGTTACGTTAGAG |
| TBP-q-R | CCTTGGGCTGTTCTAATCTG |
| SisePK2-q-F | GATGTTACCAACGTATGACGCACC |
| SisePK2-q-R | CCCTTTCCGCCTTCTCAATAACCT |
| SisCdvB-q-F | CAATGAGAAGAGGCGAAAGGCTC |
| SisCdvB-q-R | ATCATCCCCTTCTACTTGTGCCC |
| SisCdvB1-q-F | GCCATGCTCGGAAAAGATTTCC |

| | |
|---|---|
| SisCdvB1-q-R | GAAATGTAAGCGTCAAGCCTAC |
| pSeSD-F | TGGCGGTACATAGTGGTACATTAAAGTA |
| pSeSD-R | AAACCTTATGTTAAACTACGCCAGTAGG |

* The restriction sites are in bold.

**Supplementary dataset.** Summary of phosphoproteomic analysis of the ePK2 overexpression strain.
